# Tensor decomposition based feature extraction and classification to detect natural selection from genomic data

**DOI:** 10.1101/2023.03.27.527731

**Authors:** Md Ruhul Amin, Mahmudul Hasan, Sandipan Paul Arnab, Michael DeGiorgio

## Abstract

Inferences of adaptive events are important for learning about traits, such as human digestion of lactose after infancy and the rapid spread of viral variants. Early efforts toward identifying footprints of natural selection from genomic data involved development of summary statistic and likelihood methods. However, such techniques are grounded in simple patterns or theoretical models that limit the complexity of settings they can explore. Due to the renaissance in artificial intelligence, machine learning methods have taken center stage in recent efforts to detect natural selection, with strategies such as convolutional neural networks applied to images of haplotypes. Yet, limitations of such techniques include estimation of large numbers of model parameters under non-convex settings and feature identification without regard to location within an image. An alternative approach is to use tensor decomposition to extract features from multidimensional data while preserving the latent structure of the data, and to feed these features to machine learning models. Here, we adopt this framework and present a novel approach termed *T-REx*, which extracts features from images of haplotypes across sampled individuals using tensor decomposition, and then makes predictions from these features using classical machine learning methods. As a proof of concept, we explore the performance of *T-REx* on simulated neutral and selective sweep scenarios and find that it has high power and accuracy to discriminate sweeps from neutrality, robustness to common technical hurdles, and easy visualization of feature importance. Therefore, *T-REx* is a powerful addition to the toolkit for detecting adaptive processes from genomic data.

## Introduction

Natural selection refers to the evolutionary processes that differentially affect the number of offspring organisms may leave in the next generation based on the fitness of particular traits in an environment [Gillespie, 2004]. As traits will typically have some genetic basis, changes in the frequencies of traits in the population will also influence frequencies of genetic variants, or alleles, that contribute to these traits. Specifically, positive natural selection is the process by which beneficial traits increase in frequency in a population, leading to increases in the frequencies of alleles coding for the traits they contribute to, and ultimately a decrease in genetic variation at the locus under selection [Gillespie, 2004]. Because positive selection may cause particular alleles to rapidly rise in frequency in a population, through the process of genetic hitchhiking neutral genetic variants at sites nearby the selected locus will also rise to high frequency with it [Maynard Smith and Haigh, 1974, Przeworski, 2002, Kim and Nielsen, 2004, Hermisson and Pennings, 2017]. This indirect influence of positive selection on neighboring sites causes a loss of neutral genetic variation, resulting in the phenomenon coined as selective sweep [Hermisson and Pennings, 2005, Pennings and Hermisson, 2006a,b].

Inferences of such selective sweep events have been important for learning about a number of traits, such as how some human populations have evolved to digest lactose after infancy due to the advent of agriculture [Tishkoff et al., 2007b, Field et al., 2016, Ségurel and Bon, 2017, Taliun et al., 2021], the ability of organisms to survive at extreme environments such as high altitudes [Beall et al., 2010, Bigham et al., 2010, Simonson et al., 2010, Yi et al., 2010, Peng et al., 2011, Wang et al., 2011, Xu et al., 2011, Huerta-Sánchez et al., 2013, 2014, Zhang et al., 2014, Wei et al., 2016, Lindo et al., 2018, Graham and McCracken, 2019, Liu et al., 2019, Szpiech et al., 2021, Zhang et al., 2021], and the rapid spread of certain viral variants that require societies to regularly generate new drugs and vaccines [Rambaut et al., 2008, Bedford et al., 2011, Feder et al., 2016, Kim and Kim, 2016, Feder et al., 2021, Kang et al., 2021]. These important applications to human and other study systems have fueled significant interest in detecting sweeps among evolutionary, ecological, anthropological, and epidemiological researchers over the last several decades. Initial efforts toward identifying signatures of selective sweeps from genetic data were with summary statistics, which classically explored deviations from expected genetic variation under simple models of neutrality. Such approaches have been expanded in recent years, to employ diverse forms of variation, such as haplotype diversity within and among populations to increase both power to detect sweeps and robustness against confounding factors [Sabeti et al., 2002, Voight et al., 2006, Sabeti et al., 2007, Ferrer-Admetlla et al., 2014, Garud et al., 2015, Harris et al., 2018, Torres et al., 2018, Harris and DeGiorgio, 2020a, Szpiech et al., 2021]. However, with the growth in computational power and theoretical advances for modeling sweeps, complementary model-based approaches have become ever more common, as they provide a probabilistic approach for detecting sweeps and typically exhibit greater power than summary statistic approaches, provided assumptions of the underlying model fits observed data well enough [Kim and Stephan, 2002, Nielsen et al., 2005b, Chen et al., 2010, Huber et al., 2015, Vy and Kim, 2015, DeGiorgio et al., 2016, Racimo, 2016, Lee and Coop, 2017, Harris and DeGiorgio, 2020b, Setter et al., 2020, DeGiorgio and Szpiech, 2022]. Yet, these approaches still suffer in that the complexity of scenarios they can model are limited, as they are typically grounded in simple theoretical models for expected genomic variation.

Instead, due to a renaissance in artificial intelligence, machine learning methods have been at the forefront of recent efforts for detecting natural selection events from patterns in genomic variation [Schrider and Kern, 2018]. A number of approaches employ multiple summary statistics as input features, and differ in the types of summary statistics and the way at which input features are modeled [Lin et al., 2011, Schrider and Kern, 2016, Sheehan and Song, 2016, Kern and Schrider, 2018, Sugden et al., 2018, Mughal and DeGiorgio, 2019, Mughal et al., 2020, Arnab et al., 2022, Lauterbur et al., 2022]. Because the summary statistics target different patterns of genetic variation, the ensemble of such statistics can be used to provide cumulative evidence for, or against, the probability of a selective sweep producing the set of summary statistic values. Importantly though, these machine learning approaches require that hand-engineered summary statistics are chosen in advance, when they may not necessarily be the best features for discriminating among diverse evolutionary events. As a complementary strategy concurrent with the rise of deep learning [LeCun et al., 2015], convolutional neural networks [CNNs; LeCun et al., 1998] have been recently employed as a mechanism to automatically extract features and detect sweeps from raw genotypic variation [Chan et al., 2018, Flagel et al., 2019, Torada et al., 2019, Isildak et al., 2021, Gower et al., 2021]. To use CNNs as a way to extract features and detect selective sweeps, the genomic region has to be represented as images, and such approaches have matched or outperformed other statistical frameworks [Kern and Schrider, 2018, Flagel et al., 2019, Torada et al., 2019, Isildak et al., 2021].

CNNs are powerful tools that have proven useful in image classification and deep learning tasks [LeCun et al., 1998, Gu et al., 2018]. Despite their robustness, they may suffer some limitations for detecting sweeps. Because the majority of CNN architectures have at least one fully-connected dense hidden layer prior to the output layer, such models often have an enormous number of parameters [Goodfellow et al., 2016b]. The increased number of parameters generally requires larger training sets to learn their parameters, and the computational complexity of finding the optimal parameters is often high. Moreover, CNN architectures are typically agnostic with respect to where in an input image an object to detect is located, thereby ignoring important information when detecting selective sweeps, as haplotype diversity should be altered nearby a selected locus [e.g., Hermisson and Pennings, 2005, Pennings and Hermisson, 2006a,b] and support for a sweep centered on a particular genomic location should change depending on whether the altered diversity is at the center or periphery of the image. Instead, it may be useful to employ techniques that automatically extract features from images while retaining the spatial location within the image of important features, and to then use these features as input to the many powerful linear and nonlinear machine learning methods that have been developed [Hastie et al., 2009]. One such approach for extracting features from image data is tensor decomposition [Kolda and Bader, 2009].

Tensor decomposition is a class of dimensionality reduction techniques that can be applied to extract important features from data that has higher-order structure [Kolda and Bader, 2009]. Data with higher-order structure differs from typical data that is collected as a vector of feature values, as the feature values are organized in a specific manner. For example, image data has higher-order structure, as pixel (feature) values are organized into rows and columns, with pixels tending to have similar values if they have similar row-column coordinates. Traditional data analysis methods need higher-order data to be flattened into a vector for each observation before it can be analyzed. Moreover, this flattening procedure runs the risk of erasing information that might be encoded within the higher-order structure of the data. In situations where it is important to maintain the integrity of the structure of such higher-order structured data, tensor decomposition can be a useful tool for embedding this higher-order structured data in a low-dimensional space while retaining the information encoded in the original data. Tensor decomposition when applied to higher-order data can extract features, which in turn can be used for prediction tasks such as classification.

Additionally, working with high-dimensional data containing enormous numbers of features comes with an increased computational cost for a predictive model, which sometimes is referred to as “curse of dimensionality” [Bellman, 1966]. Most nonlinear methods suffer more from this curse of dimensionality than linear methods, as nonlinear methods involve a large number of parameters [Verleysen and François, 2005]. To circumvent this curse of dimensionality issue, dimensionality reduction-based [Salem and Hussein, 2019] and ensemble-based methods [Sun et al., 2020] have been developed that operate on vector representations of data, whereas tensor decomposition-based dimensionality reduction techniques are able to also retain the spatial information of features in data that have higher-order structure [Kolda and Bader, 2009].

Feature extraction is one of the foremost steps for classifying data, and tensor decomposition has emerged as an efficient approach to extract a small number of features from high-dimensional data. When extracting features from images of raw genomic data, the curse of dimensionality emerges as a problem for which traditional dimensionality reduction approaches (*e*.*g*., principal component analysis) are unideal solutions as they do not retain the spatial structure of the images. Also, many classical machine learning algorithms, such as support vector machines (SVMs), take only feature vectors as input for image data (feature matrices) must be converted into first-order tensors (vectors), which not only compromises the spatial structure of the input data but also is prone to classification errors [Liu, 2021].

In this article, we introduce a set of methods termed *T-REx*(Tensor decomposition-based Robust feature Extraction and classification) that utilize tensor decomposition for automatic feature extraction and classification of genomic image data with an aim toward distinguishing sweep footprints from neutrality. We decompose genomic data obtained from images of haplotypes using CANDECOMP/PARAFAC (CP) decomposition [Carroll and Chang, 1970b, Harshman, 1970], which is a popular model for tensor decomposition. After decomposition, the tensor is expressed as an outer product of three factors, each of which are vectors, resulting in retention of spatial structure. We feed these extracted features as input to classical linear and nonlinear classifiers to predict whether genomic regions represented as images show properties consistent with positive natural selection or neutrality. We also performed an empirical analysis using variant calls from whole-genomes of a central European population curated from the 1000 Genomes Project [The 1000 Genomes Project Consortium, 2015], in which we found novel candidate sweep genes (*e*.*g*., *MIR6874, ZNF815P, OCM*, and *SNHG17*) as well as recapitulated prior findings from the literature (*e*.*g*., *LCT, MCM6, SLC45A2*, and *EMC7*). Finally, we implemented *T-REx* as open-source software, which is available at https://github.com/RuhAm/T-REx.

## Results

The objective of *T-REx* is to automatically extract features from high-dimensional genomic data using tensor decomposition [Kolda and Bader, 2009], and to use these features to build a model to detect patterns of adaptation in genomes. To explore the efficacy of *T-REx* for detecting sweeps, we considered a diverse array of factors that can ultimately influence method power, accuracy, and robustness. We first evaluated how machine learning architecture affected accuracy and power, exploring both linear and nonlinear modeling frameworks [Hastie et al., 2009]. We then considered how the confounding effects of nonequilibrium demographic history and missing genomic segments alter relative sweep classification ability. Across all these settings, we directly compared *T-REx* to a leading sweep classifier, `ImaGene` [Torada et al., 2019], which also uses images of haplotype alignments as input. Finally, based on these simulation results, we apply the best strategy to whole-genome sequences from central European human individuals [The 1000 Genomes Project Consortium, 2015], and compare our findings to previously-reported results from the literature.

### Feature extraction and model training

To generate training and testing data for *T-REx*, we created two datasets of varying degrees of constraint associated with them. These datasets are simulated under a constant population size demographic history of 10,000 diploid individuals [Takahata, 1993, Laurent et al., 2013] with the coalescent simulator `discoal` [Kern and Schrider, 2016] using a uniform per-site per-generation mutation rate of 1.25 × 10^*−*8^ [Scally and Durbin, 2012] and per-site per-generation recombination rate of 10^*−*8^ [Payseur and Nachman, 2000] drawn from an exponential distribution and truncated at three times the mean [Schrider and Kern, 2016]. The length of the sequences was set to 1.1 megabases (Mb), and we sampled 200 haplotypes from each simulation under this setting.

In addition to these parameters, to simulate selective sweeps we introduced a beneficial mutation at the center of the simulated sequences and set the per-generation selection coefficient *s* ∈ [0.005, 0.5], which was sampled uniformly at random on a logarithmic scale. We set the initial frequency of the beneficial allele at the time of selection to be *f* ∈ [0.001, 0.1], which was also sampled uniformly at random on a logarithmic scale. This range for *f* allowed us to explore both hard and soft sweeps [Hermisson and Pennings, 2017]. The beneficial mutation became fixed *t* generations prior to sampling, and we created two datasets based on the distribution of *t* that are of varying difficulty to discriminate sweeps from neutrality. In the first dataset (denoted by `constant_1`), we set *t* = 0, and in the second more challenging dataset (denoted by `constant_2`), we draw *t* ∈ [0, 1200] uniformly at random, thereby permitting greater overlap between sweep and neutral classes. Using this protocol, we independently generated 10,000 training and 1000 test observations per class for each dataset. We developed an approach for processing haplotype alignments that may make the structure of input images easier to discern by CP decomposition. Full details of this alignment processing strategy are provided in the *Methods*.

For each dataset (`constant_1` or `constant_2`), using the `rTensor` package [Li et al., 2018], we performed a rank *R* CP tensor decomposition across a set of 20,000 training observations (10,000 per class) to obtain a low-dimensional representation of the observations in *R*-dimensional space. Using Equation 3 in the *Methods* section, we projected the 2000 (1000 per class) test observations of processed image alignments onto the *R*-dimensional subspace learned from the training set. The *CP tensor decomposition* subsection of the *Methods* provides a detailed overview of CP tensor decomposition, including learning the low-dimensional representation of the training set and projection of the test observations onto this subspace. Identifying an appropriate rank or number of components (*R*) is a key task for performing CP decomposition, yet an exact algorithm does not exist for finding the optimum *R* that gives the best approximation to the original tensor [Kolda and Bader, 2009]. Because the performances of our classifiers vary greatly across different ranks, we evaluated different values of rank *R* ∈ {50, 100, 150, 200, 250, 300} until we identified a rank that yielded excellent power and accuracy while remaining computationally efficient.

After extracting the factor matrices **A, B**, and **C** upon performing CP tensor decomposition, we fed the extracted features from the factor matrix **A** (details are provided in the *Methods*) into both classical linear (elastic net logistic regression [EN]) and nonlinear (support vector machine with a radial basis kernel [SVM] and random forest [RF]) models. We refer to these EN, SVM, and RF algorithms integrated within *T-REx* as *T-REx*(EN), *T-REx*(SVM), and *T-REx*(RF), respectively (details on training each classifier in *Methods* section). The pipeline outlining the overall procedure, from feature extraction via CP tensor decomposition to classification of genomic regions as neutral or sweep, is illustrated in Figure 1.

**Figure 1:**
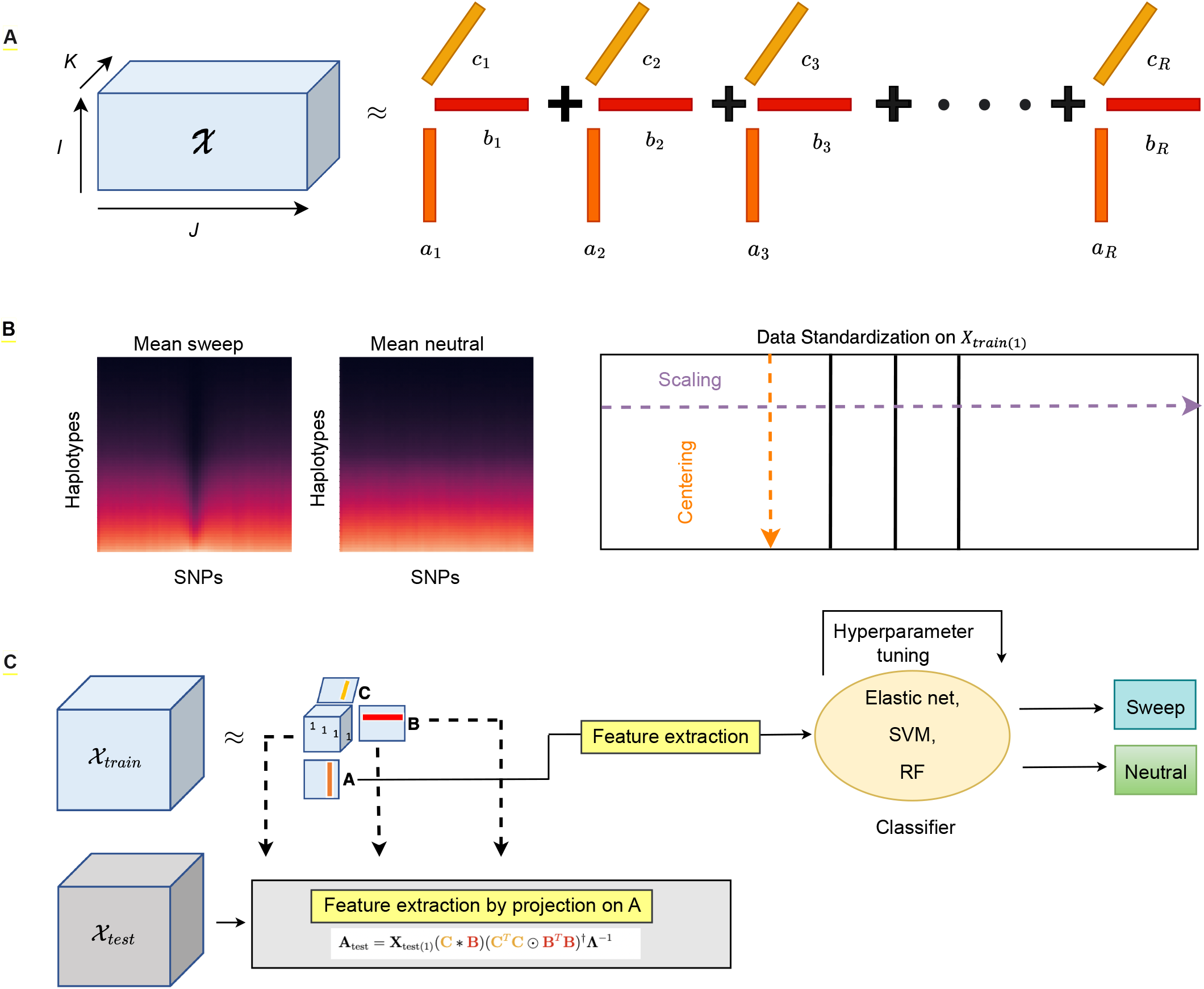
(A) CANDECOMP/PARAFAC (CP) decomposition of three-way training tensor 𝒳 ∈ ℝ^*I*×*J*×*K*^ reduces the tensor into *R* rank-one components where **a**_*r*_ ∈ ℝ^*I*^, **b**_*r*_ ∈ ℝ^*J*^, and **c**_*r*_ ∈ ℝ^*K*^ for *r* = 1, 2, …, *R*. (B) Heatmaps illustrate mean images for sweep and neutral class simulations with haplotypes along rows and SNPs along columns, with mean taken across *I*/2 training observations for each class (*I* = *N*_train_ is the total number of training observations across classes). Each cell of the image is a minor allele frequency value ranging from zero (darker colors) to one (brighter colors) representing the mean number of copies of the minor allele for the haplotype on row *j* ∈ {1, 2, …, *J* } at SNP in column *k* ∈ {1, 2, …, *K*}, where the average is taken across overlapping windows during image processing (see *Methods*). Rows are sorted from top to bottom of the image with increasing *L*_2_-norm taken across the *K* columns. Therefore, haplotypes toward the top of the image have on average a greater number of SNPs with the major allele than haplotypes toward the bottom. This sorting demonstrates that near the center of the *K* columns (where selection occurs in sweep simulations), there is a greater number of haplotypes with the major allele (darker colors) at many SNPs. The right figure in panel B illustrates the standardization process, where the mode-1 unfolded (matricized) data is centered and scaled along the columns and rows, respectively. (C) Feature extraction from the training data and the testing data is based on factor matrix **A** from the CP decomposition. For training data, the matrix **A** is obtained from CP decomposition on the training tensor 𝒳_train_, whereas the corresponding **A**_test_ factor matrix for the test dataset is obtained by projecting the test observations onto this factor learned from the training dataset. This projection is accomplished using the displayed equation, where **X**_test(1)_ is the mode-1 unfolding (matricization) of the tensor 𝒳_test_, the superscript *T* denotes transpose, the symbol ∗ denotes the Khatri-Rao product, the symbol ⊙ denotes the Hadamard (element-wise) product, the superscript † denotes the Moore-Penrose pseudoinverse, and where **Λ**^*−*1^ represents the inverse of the diagonal matrix **Λ** ∈ ℝ^*R×R*^ of scaling terms *λ*_1_, *λ*_2_, …, *λ*_*R*_. The extracted features are fed to a classifier, which outputs the class predictions.

### Power and accuracy for detecting sweeps

We first evaluate the performance of *T-REx* under the `constant_1` and `constant_2` datasets (details are provided above in the *Feature extraction and model training* subsection of the *Results*) across different CP decomposition ranks *R* ∈ {50, 100, 150, 200, 250, 300}. We selected the model resulting from the best performing rank for each of the methods based on the smallest cross-validation loss across the ranks (Figures S1 and S2). We find that across different ranks, *T-REx*(EN) has the lowest error among the three methods and *T-REx*(RF) showed lower error than *T-REx*(SVM). For the `constant_1` dataset, *T-REx*(EN) achieves an accuracy of 93.15% and maintains relatively balanced classification rates across neutral and sweep settings, with a slight, yet conservative skew toward prediction of neutrality (Figure 2). *T-REx*(SVM) and *T-REx*(RF) have lower accuracies (87.15 and 89.70%, respectively), with *T-REx*(SVM) reaching 98.2% accuracy on neutral settings (Figure 2). For the more challenging `constant_2` dataset, *T-REx*(EN) attains accuracy of 91.55% with high classification accuracies for both sweep and neutral scenarios, and with minimal mis-classification of neutral regions as sweeps. Upon a closer look at the classification rates, we find that *T-REx*(SVM) has a high accuracy of 97.0% on neutral settings, but suffers from greater sweep misclassification than *T-REx*(EN) (Figure 2). The high power displayed by the receiver operating characteristic (ROC) curves echos the high accuracy evidenced by the confusion matrices, showing that *T-REx*(EN) has high true positive rates for low false positive rates (Figure 2).

**Figure 2:**
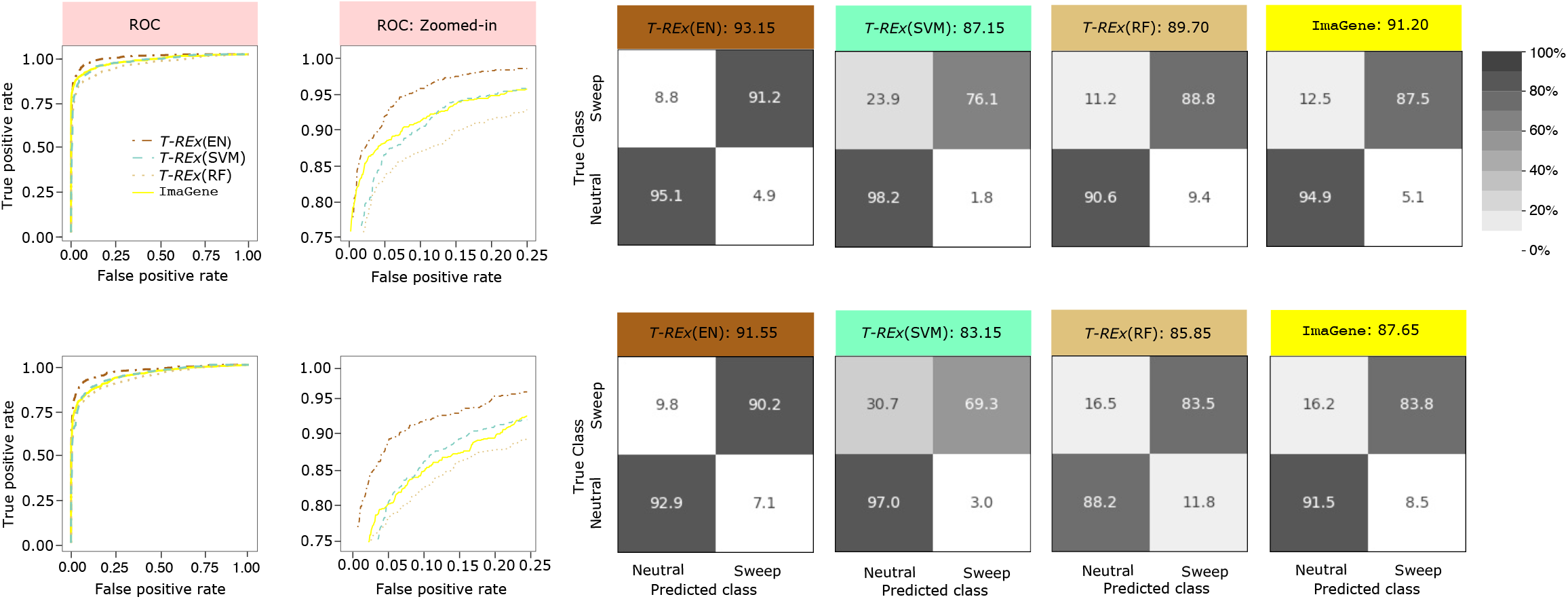
Powers and accuracies to detect sweeps for the linear *T-REx*(EN) and nonlinear *T-REx*(SVM) and *T-REx*(RF) classifiers in comparison with the CNN-based classifier `ImaGene` under a constant-size demographic history using two datasets (`constant_1` and `constant_2`) of varying difficulty. For training and testing purposes, the number of observations used for each class was 10,000 and 1000, respectively. Selective sweeps were simulated using a per-generation selection coefficient *s* ∈ [0.005, 0.5] and an initial frequency of beneficial allele at the time of selection *f* ∈ [0.001, 0.1], where both *s* and *f* were sampled uniformly at random on a logarithmic. The beneficial mutation became fixed *t* generations prior to sampling, and the distribution of *t* was set as *t* = 0 for the `constant_1` dataset (top row) and *t* ∈ [0, 1200] for the more difficult `constant_2` dataset (bottom row). For each *T-REx* method, we selected the model resulting from the best performing rank, which was chosen as the rank with the smallest cross-validation loss across the ranks *R* ∈ {50, 100, 150, 200, 250, 300}. For the `constant_1` dataset, *R* = 300, 300, and 50 were chosen for *T-REx*(EN), *T-REx*(SVM), and *T-REx*(RF), respectively, and for the `constant_2` dataset, *R* = 300 was chosen for all *T-REx* methods. Powers to detect sweeps of all four methods are compared using receiver operating characteristic (ROC) curves (first column) and ROC curves zoomed in to the upper left-hand corner with false positive rate less than 0.25 and true positive rate greater than 0.75 (second column). Classification accuracy and rates of all four methods are depicted using confusion matrices in columns three through six for *T-REx*(EN), *T-REx*(SVM), *T-REx*(RF), and `ImaGene`, respectively.

By comparing our methods to the CNN-based classifier `ImaGene` [Torada et al., 2019], we find that *T-REx*(EN) surpasses `ImaGene` in terms of power, accuracy, and classification balance on both datasets (Figure 2). However, `ImaGene` outperforms *T-REx*(SVM) in terms of power, accuracy, and classification balance whereas *T-REx*(RF) has slightly more balanced classification rates than `ImaGene`. Though all methods of *T-REx* and `ImaGene` have a skew toward predicting neutrality, `ImaGene` mistakes sweeps for neutrality more often than *T-REx*(EN) (Figure 2), which drives the lower accuracy and power of ImaGene relative to *T-REx*(EN).

### Performance under population size changes

The constant-size demographic history underlying the `constant_1` and `constant_2` datasets is an idealistic model and does not capture the fluctuations in population size often experienced by real populations [Beichman et al., 2018]. In particular, demographic scenarios, such as strong and recent population bottlenecks, which lead to an overall loss of haplotypic diversity across the genome as well as an increase in the variance of coalescence times, have been shown to generate false signatures of sweeps as well as reduce the power of sweep detection [Jensen et al., 2005]. Therefore, to investigate the performance of *T-REx* on a nonequilibrium setting with population size fluctuations and a strong, recent population bottleneck, we simulated data under a demographic history inferred [Terhorst et al., 2019] from the central European (CEU) human individuals of the 1000 Genomes Project dataset [The 1000 Genomes Project Consortium, 2015].

The distributions that selection parameters were drawn from and the number of simulated replicates per class were identical to the constant-size setting (details regarding the constant-size setting are provided in the *Feature extraction and model training* subsection of the *Results*). Analogous to the two constant-size models, we generated a dataset (denoted by `CEU_1`) where we set *t* = 0 as well as a second dataset (denoted by `CEU_2`) representing a more complicated setting where we draw *t* ∈ [0, 1200]. For each dataset, we consider an array of ranks *R* ∈ {50, 100, 150, 200, 250, 300} and compared *T-REx* to the CNN-based sweep classifier `ImaGene`.

Similar to evaluation of the two constant-size datasets, we chose the best model through cross-validation and *T-REx*(EN) generally showed the lowest error, followed by *T-REx*(RF) and *T-REx*(SVM) across different ranks. Among all the methods considered, we find that *T-REx*(EN) generally has highest accuracy and power on both the `CEU_1` and `CEU_2` datasets (Figure 3). Additionally, *T-REx*(EN) showed the lowest error in general among the three models selected from their optimal ranks. On either dataset, *T-REx*(EN) generally exhibits an increase in accuracy with increase in *R*, whereas the opposite tendency holds for *T-REx*(SVM) and *T-REx*(RF) in which their highest accuracies were attained with a small *R* value (Figures S3 and S4). This trend in the accuracy of *T-REx*(EN) with increasing *R* appears to be primarily driven by decreases in the rate of misclassifying sweeps as neutral, leading to more balanced classification rates. However, *T-REx*(EN) also achieves higher accuracy on neutral settings with increasing *R*, which is desirable as it limits false discovery of sweeps. Finally, we find, as expected, that accuracies for all methods tend to be lower for the more complex `CEU_2` dataset compared to `CEU_1` (Figure 3).

**Figure 3:**
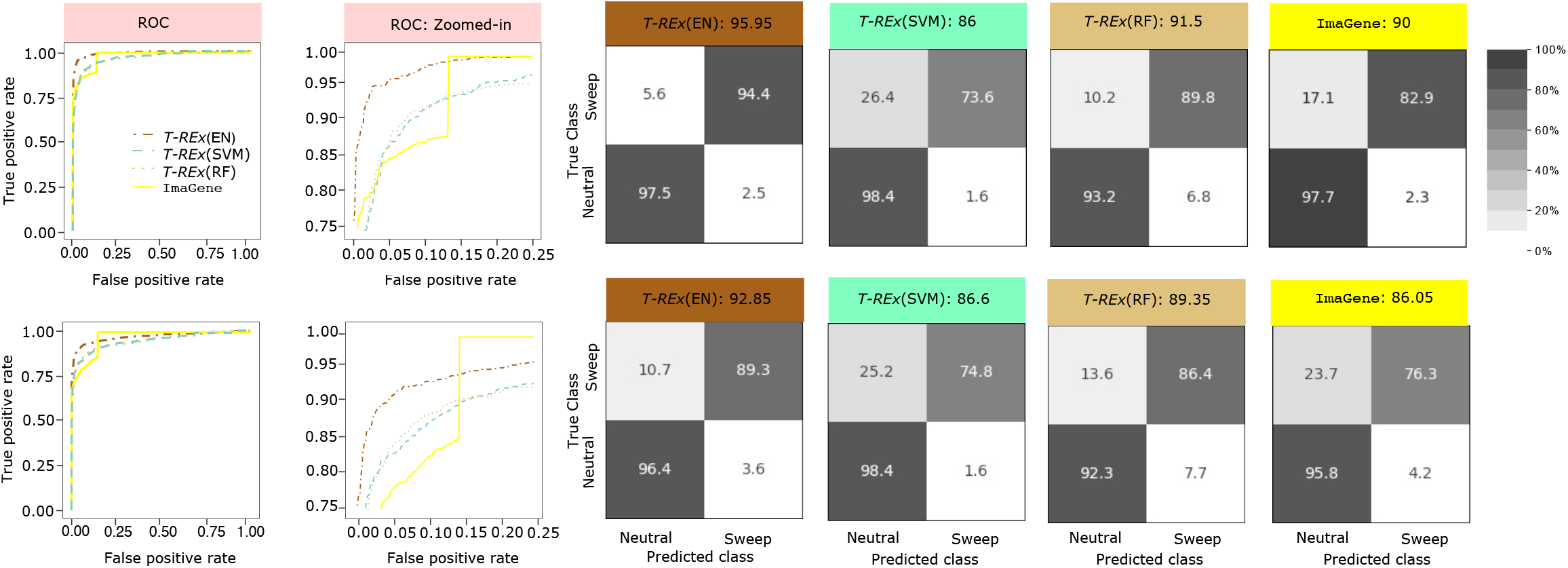
Powers and accuracies to detect sweeps for the linear *T-REx*(EN) and nonlinear *T-REx*(SVM) and *T-REx*(RF) classifiers in comparison with the CNN-based classifier `ImaGene` under a demographic history inferred from the central European human (CEU) population [Terhorst et al., 2019] history using two datasets (`CEU_1` and `CEU_2`) of varying difficulty. For training and testing purposes, the number of observations used for each class was 10,000 and 1000, respectively. Selective sweeps were simulated using a per-generation selection coefficient *s* ∈ [0.005, 0.5] and an initial frequency of beneficial allele at the time of selection *f* ∈ [0.001, 0.1], where both *s* and *f* were sampled uniformly at random on a logarithmic. The beneficial mutation became fixed *t* generations prior to sampling, and the distribution of *t* was set as *t* = 0 for the `CEU_1` dataset (top row) and *t* ∈ [0, 1200] for the more difficult `CEU_2` dataset (bottom row). For each *T-REx* method, we selected the model resulting from the best performing rank, which was chosen as the rank with the smallest cross-validation loss across the ranks *R* ∈ {50, 100, 150, 200, 250, 300}. For both the `CEU_1` and `CEU_2` dataset, *R* = 250, 50, and 50 were chosen for *T-REx*(EN), *T-REx*(SVM), and *T-REx*(RF), respectively. Powers to detect sweeps of all four methods are compared using receiver operating characteristic (ROC) curves (first column) and ROC curves zoomed in to the upper left-hand corner with false positive rate less than 0.25 and true positive rate greater than 0.75 (second column). Classification accuracy and rates of all four methods are depicted using confusion matrices in columns three through six for *T-REx*(EN), *T-REx*(SVM), *T-REx*(RF), and `ImaGene`, respectively.

In general, *T-REx*(EN) and *T-REx*(RF) outperform `ImaGene` for both the `CEU_1` and `CEU_2` datasets in terms of power and accuracy, and *T-REx*(SVM) has similar (on `CEU_2`) or worse (on `CEU 1`) accuracy compared to `ImaGene` due to it incurring higher misclassification rates of sweeps (Figure 3). Moreover, though `ImaGene` has a low miclassification rate for neutral regions, its overall accuracy suffers due to the high misclassification rate of sweeps as neutral, similar to *T-REx*(SVM). These imbalances in classification rates, however, are conservative as `ImaGene` and *T-REx*(SVM) are not prone to false discovery of sweeps. These results reiterate the strength of CP decomposition to extract features from images, even when prediction is made with a linear model (*i*.*e*., *T-REx*(EN)). The high power of *T-REx*(EN) on the two datasets reflects its strong accuracy evidenced by the confusion matrices, with high true positive rates for low false positive rates (see ROC curves in Figure 3). We note that `ImaGene` displays a spike in power at a false positive rate of about 15% for both datasets (Figure 3), which is due to approximately 19% of the predicted sweep probabilities for `ImaGene` being exactly one. The excellent classification performance of *T-REx*(EN) on a complex bottleneck setting (the `CEU_2` dataset) is promising, and so we will apply it to whole-genome data from individuals derived from the same population to scan for sweeps as a proof of concept of our prediction framework (see *Application to human genome variation* subsection of the *Results*).

### Feature maps for model interpretability

In addition to its capacity to extract features for prediction problems, CP tensor decomposition provides a low-rank representation of the original tensor, thereby allowing a mechanism for visualizing the spatial components of the images we have collected within our training tensor through factor matrices. These feature maps provide a depiction of the image characteristics that are then fed to classification models. We generated feature maps for *R* = 250 components under the `CEU_2` dataset and these feature maps reveal part of the latent structure of the tensor, with the rows and columns of these feature maps representing haplotypes and loci, respectively. Close examination of each of the components (Figures S5 to S9) reveals gradients in each of the individual feature matrices that represent the separation of features characterized by clusters of similar colors. Though some of the components show gradients in each of the individual feature matrices and clusters of similar colors where we might expect there to be signal in the haplotype alignments to discriminate between sweeps and neutrality, creating a lucid picture of the underlying features is difficult from the set of *R* = 250 images. Moreover, these feature maps only convey information about what characteristics of images were used to separate out observations from the training set, and therefore are not guaranteed to be informative about what characteristics are important for prediction.

To address this issue, we created model-informed feature maps for both datasets through a linear combination of the *R* feature maps, weighing each map by its component’s regression coefficient in the trained *T-REx*(EN) model (Figure 4). Displaying the feature maps in this fashion enables visualization of the characteristics of haplotype alignments the trained *T-REx*(EN) models place most emphasis. The pronounced red region around the center of the SNPs alludes to the expected location of lost diversity in sweeps, which the models use to distinguish sweeps from neutrality (Figure 4). A closer look at the heatmaps suggests that the models place negative weight on these features near the center of the alignment. In contrast, there is also a large dark blue region at the bottom of each heatmap, in which the models place positive emphasis to distinguish sweeps from neutrality. Differences between sweeps and neutrality in this region are expected to be due to the most recent, strongest, and hardest sweeps in our training sets (based on the procedure that we used to process haplotype alignments; see *Methods*). Another interesting observation we can discern from Figure 4 is the white, light blue, and light red shading surrounding the dark red region, signifying that *T-REx*(EN) puts little emphasis on these areas. This lack of emphasis suggests that diversity in this region provides little extra information for discriminating between sweeps and neutrality in the *T-REx*(EN) model.

**Figure 4:**
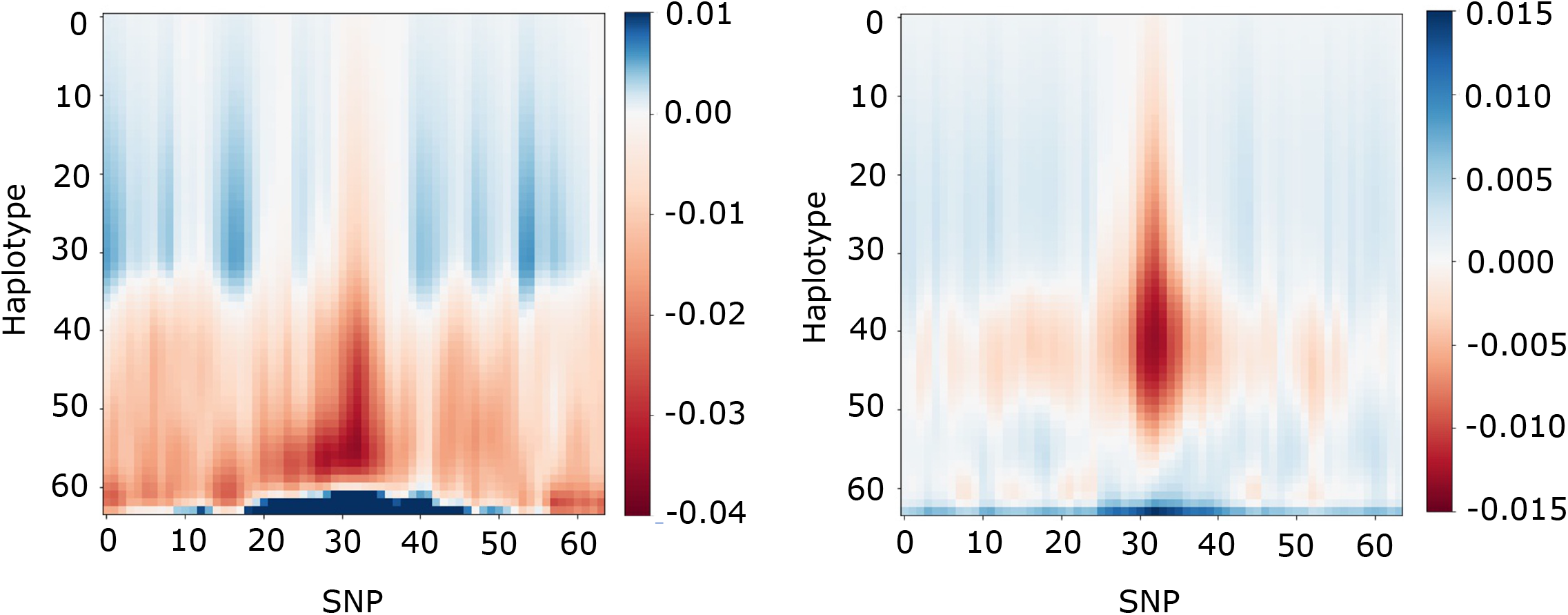
Model-informed feature maps illustrating emphasis put on different genomic regions of interest by the *T-REx*(EN) classifier trained to differentiate sweeps from neutrality under a demographic history inferred from central European (CEU) humans [Terhorst et al., 2019]. Model-informed feature maps were generated through a linear combination of the *R* feature maps (created using factor matrices **B** and **C**) from the training set, where feature map *r, r* = 1, 2, …, *R*, is weighted by the regression coefficient of component *r*(*β*_*r*_) from a trained logistic regression model with elastic net penalty. The number of components (*R*) was selected as in Figure 3 for the *T-REx*(EN) classifier, with *R* = 250 for both `CEU_1` (left panel) and `CEU_2` dataset (right panel).

### Robustness to missing data

Many genomic regions contain segments with missing SNPs, which may arise due to artifacts in the data, mapping and alignment problems, and sequencing errors. An issue that missing genomic segments poses to methods for detecting sweeps is the problem of false discovery, in which a method erroneously detects a neutrally-evolving region as a sweep [Mallick et al., 2009, Mughal and De-Giorgio, 2019]. These false signals result from the loss of SNPs in missing segments decreasing haplotypic diversity (see schematic in Figure 5), which has been shown to mislead some machine learning classifiers to call such neutral regions with confidence as sweeps if such data issues are not accounted for during model training [*e*.*g*., Kern and Schrider, 2018, Mughal and DeGiorgio, 2019]. Thus, it is important to demonstrate that *T-REx* not only has high accuracy and power to detect sweeps on idealistic data, but is robust to common technical artifacts posed by the presence of missing genomic segments.

**Figure 5:**
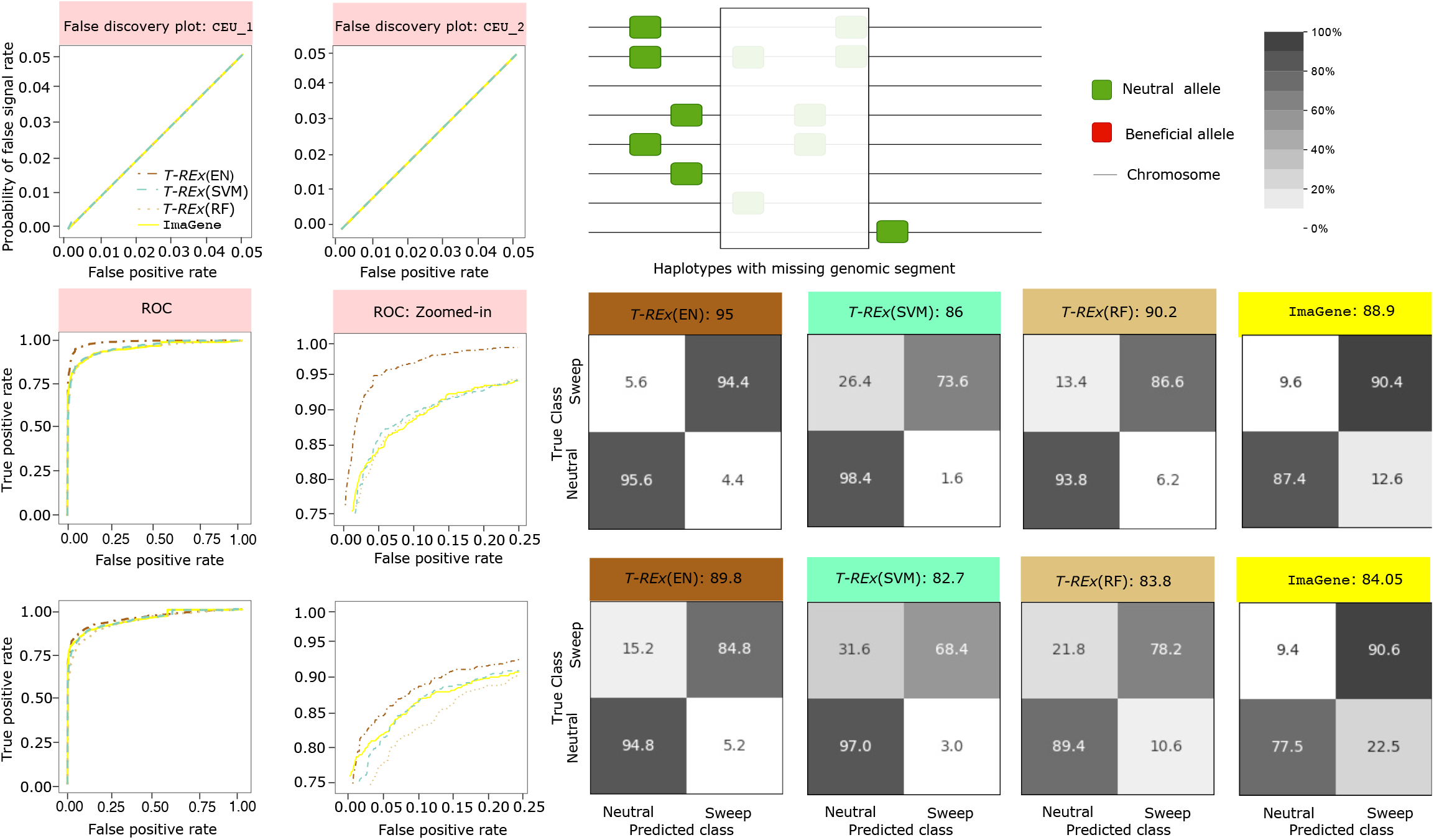
Powers, accuracies, and robustness to detect sweeps when faced with missing data for the linear *T-REx*(EN) and nonlinear *T-REx*(SVM) and *T-REx*(RF) classifiers in comparison with the CNN-based classifier `ImaGene` under a demographic history inferred from the central European human (CEU) population [Terhorst et al., 2019] history using two datasets (`CEU_1` and `CEU_2`) of varying difficulty. For training and testing purposes, the number of observations used for each class was 10,000 and 1000, respectively where 30% of the total SNPs from each test observation were removed using protocol in Mughal and DeGiorgio [2019]. (Top row) Performance of *T-REx* in comparison with `ImaGene` under missing data to ascertain whether the classifiers are robust against false discovery of sweeps, that is, erroneously detecting neutrally evolving regions as sweeps. First and second panel shows probability of false discovery of sweeps when classifying neutral genomic regions containing missing data on the `CEU_1` and `CEU_2` datasets, respectively. Third panel shows how missing genomic segment can masquerade as sweep due to apparent lack of haplotype diversity. (Middle and bottom rows) Powers to detect sweeps of all four methods are compared using receiver operating characteristic (ROC) curves (first column) and ROC curves zoomed in to the upper left-hand corner with false positive rate less than 0.25 and true positive rate greater than 0.75 (second column). Classification accuracy and rates of all four methods are depicted using confusion matrices in columns three through six for *T-REx*(EN), *T-REx*(SVM), *T-REx*(RF), and `ImaGene`, respectively. For both the `CEU_1` and `CEU_2` dataset, *R* = 250, 50, and 50 were chosen for *T-REx*(EN), *T-REx*(SVM), and *T-REx*(RF), respectively as these ranks yielded the small validation loss on nonmissing data.

The haplotype images used for training and testing sets so far have assumed no missing data, and so we seek to examine the effectiveness of our methods when test data have missing segments that may ultimately reduce observed haplotypic variation. To this end, we followed the protocol in Mughal and DeGiorgio [2019] by removing 30% of the SNPs from each test replicate to evaluate the impact of missing data on method accuracy, power, and robustness. The removal of 30% of the SNPs is accomplished in 10 non-intersecting chunks, each accounting for roughly three percent of the total SNPs in the replicate, and with starting position of each chunk chosen uniformly at random. In cases of overlap with previously-drawn missing chunks, a new starting location for the current chunk is redrawn.

Using *T-REx* models trained with non-missing data and assuming the rank *R* of CP decomposition that gave each method (*T-REx*(EN), *T-REx*(SVM), and *T-REx*(RF)) their smallest cross-validation loss, we find that on both the `CEU_1` and `CEU_2` datasets *T-REx*(EN) continues to show greater power and accuracy compared to competing approaches (center and bottom rows in Figure 5). Specifically, for both the `CEU_1` and `CEU_2` datasets, *T-REx*(EN) outperforms `ImaGene` with a margin of around 6% in terms of accuracy (center and bottom rows in Figure 5). Moreover, under both datasets, `ImaGene` is more prone to false discovery of sweeps than *T-REx*(EN), as it displays a skew toward falsely classifying neutrally-evolving regions as sweeps. In the case of the `CEU_1` dataset, *T-REx*(RF) marginally outperforms `ImaGene` in terms of accuracy. However, the accuracy of *T-REx*(SVM) suffers, as 26.4% of sweeps are misclassified (center row in Figure 5). In contrast, on the `CEU_2` dataset, `ImaGene` outperforms both *T-REx*(SVM) and *T-REx*(RF) in terms of accuracy, but falsely classifies 22.5% of the neutral observations as sweeps. This result illustrates that when presented with data containing missing genomic segments, the CNN-based classifier `ImaGene` may mistake the reduced haplotypic diversity as a sweep footprint. We expand upon this issue in the *Discussion* section, and detail procedures that can be taken to alleviate the issue of missing segments [*e*.*g*., Kern and Schrider, 2018].

To further evaluate whether *T-REx* is robust to false discovery of sweeps in neutral regions with missing data, we compute the proportion of false signals, based on the distribution of sweep probabilities of neutral replicates with missing segments, as a function of false positive rate, based on the distribution of sweep probabilities of neutral replicates without missing segments. For this purpose, we generated an additional 1000 neutral replicates each having 30% missing SNPs so that these two distributions were generated from independent neutral replicates. Sweep classifiers that are robust to neutral missing segments will have the curve relating the proportion of false signals (on the *y*-axis) as a function of the false positive rate (on the *x*-axis) fall on or below the *y* = *x* line. Our results show that for both variations of the simulated CEU dataset, curves for all tested methods fall on the *y* = *x* line, considering relevant false positive rates between zero and five percent (top row in Figure 5). We therefore conclude that all methods considered here are robust to false discovery of sweeps due to missing data when conditioning on reasonable false positive rates.

### Application to human genome variation

In addition to evaluating the performance of *T-REx* under simulated scenarios, we also embarked on an empirical application to whole-genome variant calls from a European human population as a proof of concept (details regarding processing of the empirical data are provided in the *Application to empirical data* subsection of the *Methods*). Using the identical protocol as in our assessment of model performance, we trained *T-REx*(EN) on 10,000 simulated replicates per class with parameters identical to those that generated the `CEU_2` dataset, with the exception of sampling 198 haplotypes per simulation to match the 99 diploid individuals sampled for the CEU population of the 1000 Genomes Project dataset [The 1000 Genomes Project Consortium, 2015]. We opted to apply *T-REx*(EN) for our empirical analysis, as it emerged as the best performing model among the three *T-REx* methods evaluated across a range of simulated settings.

To uncover candidate genes that show evidence of sweep signatures, we evaluated whether each gene harbored a high predicted sweep probability and a sweep probability peak, observed by computing a moving average computed as the mean of sweep probabilities at 11 contiguous genomic windows. This 11-window mean approach provides a smoothed representation of the probabilities and helps us observe the underlying trend of probability as a function of genomic position. We identified 17 regions from eight autosomes displaying pronounced peaks in predicted sweep probability, which we list together with associated genes in Table 1 and depict within Figures S10 and S11. In particular, we found candidate genes that have been supported by previous studies (*e*.*g*., *LCT, MCM6, SLC45A2*, and *EMC7*; Bersaglieri et al. [2004a], Oleksyk et al. [2010], López et al. [2014], Racimo [2016]) as well as novel candidates (*e*.*g*., *MIR6874, ZNF815P, OCM*, and *SNHG17*).

**Table 1:** Autosomal regions showing high predicted sweep probability in the CEU population as predicted by *T-REx*(EN).

| Chromosome | Start | Stop | Genes |
| --- | --- | --- | --- |
| 1 | 115,397,483 | 116,311,335 | *SYCP1, CASQ2* |
| 1 | 37,940,044 | 38,422,646 | *SF3A3, MIR4255* |
| 2 | 136,545,419 | 136,634,013 | *LCT, MCM6* |
| 5 | 33,936,490 | 33,984,798 | *SLC45A2* |
| 6 | 29,640,259 | 30,594,169 | *HLA-F, HLA-F-AS1, IFITM4P, HCG4, HLA-V, HLA-G, HLA-H, HCG4B, HLA-A, HCG9* |
| 7 | 5,751,470 | 6,369,041 | *MIR6874, ZNF815P, OCM, CCZ1, RSPH10B* |
| 7 | 27,132,611 | 27,287,449 | *HOXA1, HOXA2, HOXA3, HOXA9, HOXA10, HOXA-AS2, HOXA-AS3* |
| 10 | 15,253,641 | 15,761,921 | *FAM171A1, ITGA8* |
| 15 | 76,507,693 | 77,474,268 | *ETFA, TISL2, TYRO3P, SCAPER, RCN2, MIR3713, TSPAN3* |
| 15 | 34,376,217 | 34,649,936 | *EMC7, PGBD4, KANTBL1, EMC4, SLC12A6, NUTM1* |
| 15 | 38,988,798 | 41,248,710 | *LINC02694, C15orf54, RMDN3, GCHFR, DNAJC17, C15orf62, ZFYVE19, PPP1R14D, SPIT1-AS1, SPIT1, VPS18, LOC105370943, DLL4, CHAC1* |
| 17 | 29,861,900 | 29,902,540 | *MIR4724, MIR193A, MIR4725, MIR365B* |
| 17 | 41,453,295 | 41,864,988 | *LINC00910, ARL4D, MIR2117HG, DHX8, MEOX1, SOST, DUSP3, CFAP97D1* |
| 20 | 37,230,451 | 37,401,163 | *ARHGAP40, SLC32A1, ACTR5* |
| 20 | 50,700,549 | 51,266,965 | *ZFP64, LINC01524* |
| 20 | 37,049,234 | 37,358,015 | *SNHG17, SNORA71B, SNORA71C, SNORA71D, SNORA71E, SNORA60, RALGAPB, ADIG, ARHGAP40, SLC32A1* |
| 22 | 40,139,048 | 40,439,538 | *ENTHD1, GRAP2, FAM83F* |

#### Sweep candidates supported by the literature

On chromosome 2, we find a peak surrounding the region containing the genes *LCT* and *MCM6* (Figure 6A). In particular, we see a clear peak that reaches an 11-window mean sweep probability close to one near *LCT* and *MCM6* and decays in value with distance from these genes. This trend of reduction in sweep probability with distance from a putative adaptive locus is consistent with the footprint of a selective sweep, and is due to the action of recombination breaking down linkage disequilibrium and shaping haplotypic diversity across the chromosome [Slatkin, 2008]. *LCT* encodes the enzyme lactase that aids in lactose digestion in humans, and is a strong selection candidate, especially across European populations as the ability to digest lactose persists into adulthood within individuals of European ancestry [Scrimshaw and Murray, 1988]. This lactose tolerance is an outcome of positive selection owing to the advent of farming that resulted in an infusion of milk as part of regular consumption within particular cultures in the last 1,000 years [Sabeti et al., 2006]. Moreover, near *LCT*, we also detect the gene *MCM6* with high confidence, which has been hypothesized to have undergone positive selection by previous studies [*e*.*g*., Shatin, 1968, Harris and Meyer, 2004, Bersaglieri et al., 2004b, Nielsen et al., 2005a, Sabeti et al., 2007, Tishkoff et al., 2007a, Itan et al., 2009, Ingram et al., 2009, Schlebusch et al., 2012, Fan et al., 2016, Cheng et al., 2017]. *MCM6* contains two introns, one of which harbors an enhancer that acts as a regulatory mechanism for *LCT* and therefore may contribute to lactase persistence and have been positively selected in the past [Anguita-Ruiz et al., 2020].

**Figure 6:**
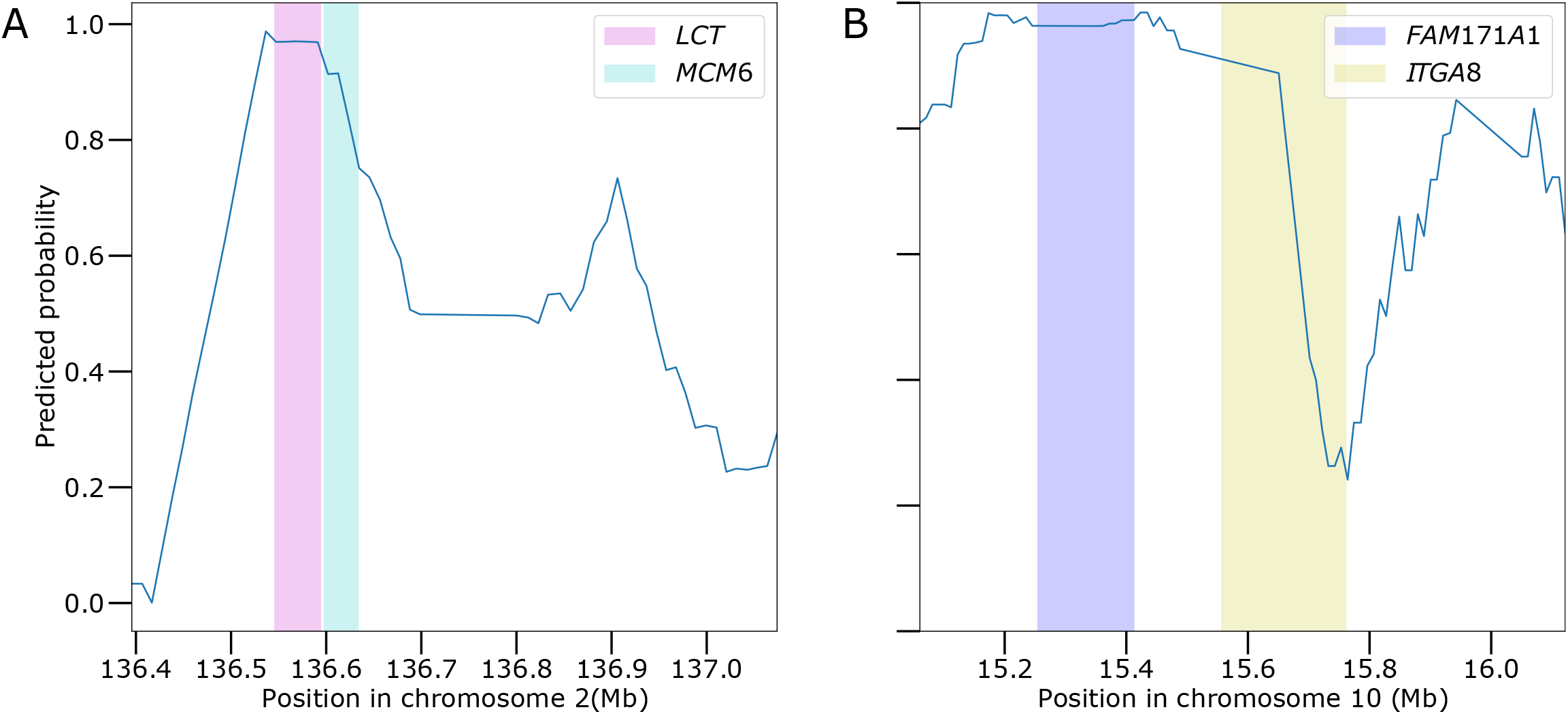
Detection of two example genomic regions containing sweep signatures within the CEU population of the 1000 Genomes Project dataset. *T-REx*(EN) predicted sweep probabilities as a function of chromosomal position surrounding the *LCT* and *MCM6* regions on chromosome 2 (panel A) and the *FAM171A1* region on chromosome 10 (panel B). The probabilities are calculated as 11-window moving averages, computed with five windows before and five windows after a given central window. The genomic intervals containing each gene are shaded using colors in accordance with the order of their appearance in the labels.

The region surrounding *LCT* and *MCM6* represents a positive control, as we expect most sweep detection methods to uncover this region with high confidence. We next went on to probe for other well-studied candidates of natural selection, and found evidence for sweeps in the major histocompatibility complex (MHC) region on chromosome 6 (Figure S10E). Specifically, *T-REx* identified high sweep support for the genes *HLA-H, HCG4B, HLA-A*, and *HCG9*, which had 11-window mean sweep probabilities close to one. Other candidate genes with moderate support in the region include *HLA-F, HLA-F-AS1, IFITM4P, HCG4, HLA-V*, and *HLA-G*, with 11-window mean sweep probabilities ranging from 0.65 to 0.81. Many genes located in the MHC region code for proteins that aid in pathogen immune defense through peptide binding [Mladkova and Kiryluk, 2017]. Loci in such genes tend to be highly polymorphic, and have long been hypothesized as evolving under balancing selection, likely due to the evolution of the host in the face of pathogens and parasites [Lederberg, 1999, Bernatchez and Landry, 2003]. The high structural variation coupled with extreme polymorphism in this region makes variant calling difficult [Stipoljev et al., 2020], and potentially poor genotype calls may have contributed toward the ambiguity in detecting sweeps in this region. Though often having different genomic footprints to positive selection, balancing selection is a clear deviation from neutrality and *T-REx* was able to identify the lack of neutrality at the MHC region. The classification of this region as positive selection by *T-REx* may be partially due to its extreme levels of linkage disequilibrium [Stipoljev et al., 2020], consistent with expectations of sweeps. However, our results are also consistent with prior studies, which have found evidence for sweep-like signals at the MHC region in humans [*e*.*g*., Campbell et al., 2019, **?**].

The gene *SLC45A2* (Figure S10D) on chromosome 5 has moderate sweep support with 11-window mean sweep probabilities around 0.75. This gene encodes a protein that plays a crucial role in melanin synthesis that affects skin pigmentation in humans [López et al., 2014]. The frequencies of alleles in this gene that are associated with pigmentation in Europeans demonstrate a latitudinal cline across Europe, resulting in lighter skin pigmentation in northern Europe [Norton et al., 2007]. Patterns of variation mimicking footprints of positive selection near *SLC45A2* in European humans are supported by numerous studies [*e*.*g*., Hider et al., 2013, Laayouni et al., 2014, López et al., 2014, MGoodwin et al., 2017, Wilde et al., 2014, MGoodwin et al., 2017].

Further investigation into the regions with high sweep support revealed *EMC7* (Figure S11B), which codes for a protein that is an important part of the endoplasmic reticulum membrane and acts as a molecular tether enabling the transport of viruses between different cellular compartments [Bagchi et al., 2020]. *T-REx* detects *EMC7* with an 11-window mean sweep probability of 0.86, which has prior support for positive selection [Racimo, 2016]. Moreover, with 11-window mean sweep probabilities ranging from 0.95 to 0.99, *T-REx* captured the genomic region containing the protein coding gene *SF3A3* (Figure S10B) on chromosome 1. García-Cárdenas [2022] demonstrated a possible connection between *SF3A3* and breast cancer and a network of cancer driving genes. Though potentially associated with the harmful disorder of cancer in contemporary environments, Racimo et al. [2014] also suggested that *SF3A3* may have been subjected to past positive selection.

#### Novel sweep candidates

In addition to these previously-identified sweep candidates, we uncovered a number of novel candidates. On chromosome 1, we found *SYCP1* (Figure S10A) as a possible sweep candidate with 11-window mean sweep probabilities reaching 0.88. This protein coding gene is part of the synaptonemal complex, which is a protein structure that forms between homologous chromosomes [Seo et al., 2016]. Hosoya and Miyagawa [2021] highlight that some of the proteins coded by *SYCP1* are abnormally expressed in 13 different cancer tissues, including breast and stomach cancer, and acute myelogenous lukemia. Moreover, mutations in *SYCP1* have been associated with male infertility [Nabi et al., 2022].

On chromosome 7, we found candidate genes belonging to the HOXA-family that exhibit high sweep support with 11-window mean sweep probabilities ranging from 0.80 to 0.95 (Figure S10H). HOXA genes are part of the homeobox cluster, which encode proteins that play an important part in early development of humans by performing embryo segmentation [Shah N, 2010], and it has been suggested that HOXA-family genes are involved in the inception and development of human cancers [Ge et al., 2021]. Specifically, *HOXA9* (Figure S10H) is responsible for the pathogenesis of acute myelogenous leukemia, which is a cancer of the bloods and bones [Chen et al., 2019].

Additionally, *SHNG17* on chromosome 20 (Figure S11H) has high sweep support with 11-window mean sweep probabilities reaching 0.95. *SHNG17* is known to be an important factor behind gastric cancer in humans, as it is upregulated in gastric cancer tissues [Zhang et al., 2019]. Furthermore, on chromosome 10, we identified a strong peak with high sweep support at the protein coding gene *FAM171A1* (Figure S10I), which is also associated with breast cancer survival and plays an important role in immune system regulation [Parada et al., 2017]. Among our highlighted novel candidates, as well as those that are previously-identified (*SF3A3*), there is an intriguing connection between these sweep candidates and cancer proliferation and suppression. This pattern of selective sweeps at genes related to cancer was also found by other studies that developed machine learning approaches for detecting sweep [*e*.*g*., Lou et al., 2014, Schrider and Kern, 2017, 2018, Mughal et al., 2020, Arnab et al., 2022]. Detection of cancer related genes by *T-REx* as well as methods from previous studies, provides an interesting pattern that many past positively-selected genes may drive current carcinogenesis in humans.

## Discussion

In this article, we have introduced a tensor decomposition-based feature extraction and classification method termed *T-REx* that is able to differentiate sweeps from neutrality with a high degree of power and accuracy. Specifically, we found that our linear model (*T-REx*(EN)) demonstrated overall superior performance to the nonlinear models (*T-REx*(SVM) and *T-REx*(RF)) across an array of different settings, including demographic history, positive selection regime, and technical artifacts due to missing genomic segments (Figures 2, 3, and 5). Moreover, in addition to its high power and accuracy to detect sweeps, this modeling framework facilitated easy interpretation of the fitted model by providing feature maps for visualization, which convey the particular location in the haplotype alignments that the models place emphasis when discriminating sweeps from neutrality (Figure 4).

From our experiments, an unexpected observation was that the linear *T-REx*(EN) model had higher power and accuracy than the nonlinear *T-REx*(SVM) and *T-REx*(RF) models (Figures 2 and 3). It is possible that the linear model performs better here because it yields a better decision boundary between the neutral and sweep classes. However, it is more likely that other factors have played a more critical role in leading *T-REx*(EN) to have the best performance. First, the *R* components resulting from the CP tensor decomposition are not required to be independent, and may, in fact, be highly correlated [Kolda and Bader, 2009]. The elastic net regularization employed by *T-REx*(EN) has both *L*_1_- and *L*_2_-norm penalties, which are both meant to handle correlated features [Hastie et al., 2009]. In particular, the *L*_2_-norm penalty reduces the effective number of features in the model, but encourages a dense model by ensuring that all features remain included in the fitted model [Hastie et al., 2009]. In contrast, the *L*_1_-norm penalty encourages a sparse model by emphasizing fewer features and selecting out those that are redundant or irrelevant for prediction [Hastie et al., 2009]. Therefore, the *L*_1_-norm penalty employed by *T-REx*(EN) method is particularly useful in reducing the overall dimension of the input data by removing irrelevant and redundant features. This hypothesis is supported by the fact that *T-REx*(EN) tends to have non-decreasing power and accuracy with increasing *R* (Figures S3 and S4). Second, though *T-REx*(SVM) also has an *L*_2_-norm penalty [Hastie et al., 2009], this penalty does not encourage sparsity in the set of input features like the *L*_1_-norm penalty. Moreover, we employ the radial basis kernel within the *T-REx*(SVM) classifier, which requires a distance be taken between observations, and distances in high-dimensional space may not behave well due to the curse of dimensionality [Verleysen and François, 2005]. This hypothesis related to the curse of dimensionality is supported by power and accuracy of *T-REx*(SVM) tending to diminish with increasing *R*, and hence has decreasing performance with increasing numbers of input features (Figures S3 and S4).

To put forth a better perspective on the utility of the haplotype alignment processing method *T-REx* uses, we experimented with another protocol for processing haplotype alignments, which is similar to that of Torada et al. [2019]. As in Torada et al. [2019], we sorted the haplotypes along the entire 1.1 Mb genomic region, which is in contrast to the alignment processing method employed by *T-REx*, where haplotypes were sorted in a sliding window. This key difference between these two protocols may be an important factor behind the decreased false discovery of sweeps by *T-REx*(EN) (compare Figure S12 to Figure S1 and Figure S13 to Figure S2). Overall, our experiments under the constant-size demographic history across different ranks (compare Figure S12 to Figure S1 and Figure S13 to Figure S2) show that our unique alignment processing method has a distinct advantage in terms of downstream classification accuracy and power over another contemporary approach for processing haplotype alignments. If `ImaGene` adopted this local alignment processing approach, then it would have potentially resulted in performance that is more close to that exhibited by *T-REx*. Another factor that has likely impacted the performance of `ImaGene` in our study is that it is CNN-based, and CNNs typically require large training sets to achieve optimal performance [Luo et al., 2018]. In the original `ImaGene` article, Torada et al. [2019] employed 50,000 observations per class for training. In contrast, we used 10,000 observations per class for comparison purposes with *T-REx*, which may have influenced the results shown by `ImaGene`. Moreover, a key distinction between `ImaGene` and *T-REx* is that `ImaGene` uses larger resized 128 × 128-dimensional images as input, which have the potential for reduced robustness to noise compared to *T-REx*, as more noise is averaged out with its smaller 64 × 64-dimensional input images.

When analyzing modern genomic data, it is common to encounter regions with missing segments due to artifacts or sequence alignment problems, making it critical that machine learning tools remain robust to the challenge such missing data poses. In our tests with missing segments, we found that *T-REx*(EN) was fairly robust, but `ImaGene` was deleteriously affected by an increase in the misclassifcation rate of neutral regions—though for reasonable false positive rates, `ImaGene` was also robust (Figure 5). An avenue to alleviate this problem is to train classifiers with missing random segments [Kern and Schrider, 2018], which allows classifiers to learn the underlying patterns associated with missing data. Randomly removing chunks from alignments in non-overlapping windows from the training data before training classifiers has been shown to offset the deleterious effects of such missing data [Mughal and DeGiorgio, 2019, Mughal et al., 2020]. Also, filling in missing values in test data through genotype imputation [*e*.*g*., Li et al., 2010, Moritz and Bartz-Beielstein, 2017, Davies et al., 2021, Browning et al., 2021] may be another direction to combat the problem of missing data. Classifiers that are fed test data after imputing the missing values tend to be robust when faced with missing data in genomes and may achieve better prediction accuracy [Sarkar et al., 2021].

We have implemented *T-REx* as a binary classifier to differentiate sweeps from neutrality, but this modeling strategy can also be employed for broader classification problems in evolutionary genomics. For example, using mutliclass extensions to the machine learning models discussed here, the *T-REx* framework could accommodate classifiers for jointly discriminating among other evolutionary processes, such as balancing selection and adaptive introgression, in addition to neutrality and sweeps from *de novo* mutations or standing variation. To illustrate, two-dimensional representations of genomic data have been employed in multiclass models for robustly determining whether a genomic region is neutrally-evolving or has undergone a hard or soft sweep [Kern and Schrider, 2018], as well as been shown to improve discrimination of adaptive introgression from sweeps and neutrality [Mughal et al., 2020]. Moreover, Gower et al. [2021] employed images of sorted haplotype alignments as input to a CNN with the aim to detect adaptive introgression—a setting that Mughal et al. [2020] still had trouble with based on two-dimensional images derived from hand-engineered population-genetic summary statistics. Indeed, Isildak et al. [2021] showed that CNNs applied to extract features from images of haplotype alignments outperformed feed-forward neural networks applied to hand-engineered population-genetic features in discriminating between recent balancing selection and incomplete sweeps, which are two evolutionary settings that can yield similar distributions of haplotype variation and are thus difficult to tease apart. These examples highlight the promise that automatic feature extraction from image representations of haplotypic variation has for probing genomes for diverse forms of natural selection.

Throughout this article, we have explored the problem of identifying natural selection as a classification task. However, the machine learning models employed by *T-REx* are flexible, and changing from a qualitative to a quantitative output would shift the problem from a classification to a regression problem. By using a regression framework, *T-REx* could predict underlying sweep parameters, such as selection strength, frequency of the selected allele when it became beneficial, and time at which a sweep completed [Mughal and DeGiorgio, 2019]. Moreover, as in Flagel et al. [2019], framing the prediction problem as regression would allow for estimation of key demographic quantities, such as the timing and magnitude of population size changes, as well as genetic parameters, such as recombination rate. Hence, tensor decomposition represents a complementary tool for tackling an array of inference problems within population genomics that CNNs have already been demonstrated to be highly effective.

Important limitations of *T-REx* are the runtime and memory-usage associated with larger training sets (*N*) and higher ranks (*R*). In our experiments, we found that tensor decomposition took substantially greater time and memory even for modest increases in *R*. Downsampling each observation to a 64 × 64-dimensional matrix helped in reducing the complexity, and also likely aided in robustness of our models by averaging some of the noise in the input images. Moreover, we have been concerned with three-way tensors only, but if we were to consider increasing the number of dimensions, it would render the process computationally costlier than a three-way case, as the number of elements in the tensor would increase exponentially with each added dimension [Kruppa, 2017]. Also, the alternating least squares algorithm (see *Methods* section) for learning the factors matrices for CP tensor decomposition will need to find the factor matrices associated with each added dimension. For example, if we were to include ancient DNA data sampled over time as the fourth dimension in our existing pipeline, then it would be a four-way tensor where we would have an extra factor matrix **D**, which the ALS algorithm has to estimate through iteration and will incur greater runtime before reaching convergence.

We have focused on CP tensor decomposition [Hitchcock, 1927, Harshman, 1970]. However, other algorithms for decomposing tensors exist, each with their own advantages and disadvantages relative to CP decomposition. Examples are multilinear principal component analysis (MPCA) [Lu et al., 2008], Tucker decomposition [Tucker, 1966], higher-order singular value decomposition (HOSVD) [Lathauwer et al., 2000], and tensor train (TT) decomposition [Oseledets, 2011], which are widely-used alternative approaches for performing tensor decomposition [*e*.*g*., Sidiropoulos et al., 2017, Yuwang et al., 2019]. Methods such as CP decomposition, MPCA, HOSVD, and TT are closely related to Tucker decomposition [Zare et al., 2018, Yuwang et al., 2019] in their working procedures, which is based on finding the linear combination of outer products of vectors. Among these different techniques, Tucker decomposition [Tucker, 1966] is the most similar in operation to CP decomposition, as it also hinges on the idea of using alternating least squares to estimate a core tensor and factor matrices, though the core tensor produced by Tucker decomposition is not necessarily diagonal like the one CP decomposition outputs [Yuwang et al., 2019] and the ranks of the factor matrices are not constrained to be identical. Despite their similarities, CP decomposition is able to produce unique solutions unlike Tucker decomposition, where factor matrices change as the core tensor is changed [Kim et al., 2014, Zare et al., 2018]. Also, the rank-one factors generated by Tucker decomposition are orthonormal, which is not the case for CP tensor decomposition [Kim et al., 2014].

The *T-REx* methodology introduced here represents complementary approach to CNNs for automatic feature extraction of haplotype alignment images. This framework is flexible, as it permits learned features to be used in both linear and advanced nonlinear models, and can be extended into multiclass and quantitative prediction problems within evolutionary genomics. Moreover, we demonstrated that *T-REx* has an edge over a current leading CNN-based architecture in terms of power and accuracy, partially due to its unique alignment processing strategy for easier feature detection. Moreover, *T-REx* identified previously hypothesized and novel candidate sweeps in our empirical application, highlighting its efficacy in practice. Despite the promising performance metrics of *T-REx*, computation time of *T-REx* increases with increasingly higher ranks and sample sizes. However, excellent power and accuracy were achieved for modest numbers of features and training set sizes, and so we do not see this as a major hurdle for *T-REx*. Given the rapidly-changing landscape of computational approaches for learning about and uncovering evolutionary mechanisms, *T-REx* provides a bridge between modern methodologies for feature extraction and well-established classical machine learning prediction techniques.

## Methods

### CP tensor decomposition

Consider a tensor 𝒳 ∈ ℝ^*I×J ×K*^ of order three, where the first dimension will collect *I* observations of two-dimensional images each with *J* × *K* pixel values. The idea behind CP tensor decomposition is to express such a tensor as a sum of *R* tensors, where each of these tensors is expressed as the outer product of three rank one tensors. That is, we wish to estimate 𝒳 as

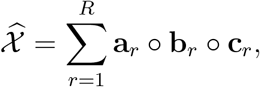

where the symbol denotes the outer product and where **a**_*r*_ ∈ ℝ^*I*^, **b**_*r*_ ∈ ℝ^*J*^, and **c**_*r*_ ∈ ℝ^*K*^ such that 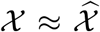. For our setting, *I* will represent the number of training observations, *J* a proxy for the number of haplotypes, and *K* a proxy for the number of loci (see *Alignment processing* subection of the *Methods* for details). Because we are working with tensors of order three, which is a higher-order tensor, we have column, row, and tube *Fibers*, which are respectively termed mode-1, mode-2, and mode-3 of the tensor.

#### Preprocessing tensors

Prior to application of CP decomposition, we need to preprocess the input tensors through centering and scaling operations. Because the data are represented as a three-way tensor, preprocessing is different from conventional methods [Kolda and Bader, 2009]. Let value *x*_*ijk*_ denote elements *i, j*, and *k* respectively for the first, second, and third dimensions of the tensor 𝒳 ∈ ℝ^*I*×*J*×*K*^. This tensor is centered as

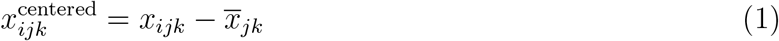

where

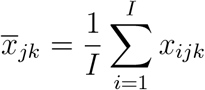

is the sample mean across the *I* training observations. Here the index *i* is related to the first mode, so it runs from 1 to *I*. Similarly index *j* runs from 1 to *J* and index *k* runs from 1 to *K*. This kind of centering is called single centering across the first mode [Bro, 1997], and causes the mean of each pixel of an image to be zero across the training samples. We could have centered on multiple modes simultaneously, which is called double or triple centering depending on the number of modes on which to simultaneously center. However, centering one mode at a time is appropriate for CP decomposition, as any other kind of centering would destroy the multilinear properties of the data [Bro, 1997]

In addition to centering, scaling should be performed on only one mode at a time, and we we have chosen to scale in the first mode for our application [Kolda and Bader, 2009]. Scaling is performed as

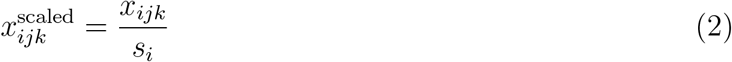

where

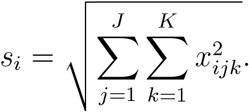

This kind of scaling ensures that the overall intensity of values across pixels in an image are identical for each training sample. The order of scaling and centering is not arbitrary, as the operations are not commutative [Kolda and Bader, 2009]. Centering across a particular mode after scaling disturbs scaling across all modes. On the other hand, scaling across a particular mode after centering destroys centering across that mode. For these reasons, the order of centering and scaling is important. Centering is performed after scaling so that the scaled mode variance is not exactly one, but any large differences across the mode are mostly equalized [Kolda and Bader, 2009]. Centering is then performed, which ensures that the mode to be centered has a mean of zero.

#### Computing the CP decomposition

After performing tensor decomposition on the training tensor 𝒳 ∈ ℝ^*I*×*J*×*K*^ to obtain a rank *R* CP decomposition, we obtain the three factor matrices

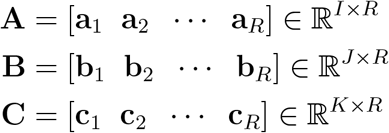

which yield an approximation of the tensor through the outer product

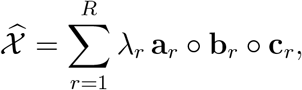

where *λ*_*r*_, *r* = 1, 2, …, *R*, scales the *r*th tensor to have unit norm. From the factor matrices **B** and **C**, we can depict the features extracted by component *r* of the CP model from the training data with the expression **b**_*r*_ ○ **c**_*r*_ ∈ ℝ^*J*×*R*^ [Papastergiou et al., 2018].

The key algorithm behind computing the CP decomposition is alternating least squares [Carroll and Chang, 1970a], which is a minimization algorithm. For a tensor of order three, given a rank *R* to approximate the training tensor (𝒳), alternating least squares fixes two of the factor matrices while solving for the remaining factor matrix that minimizes the sum of the squared differences in the elements of the estimated tensor 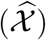 and the training tensor. For example, if factor matrices **B** and **C** are fixed, then we seek to find **A** that has this minimal sum of squared errors.

Denote the best factor matrices **A, B**, and **C** at iteration *t* ∈ {0, 1, 2, … } of the alternating least squares algorithm by **A**^(*t*)^, **B**^(*t*)^, and **C**^(*t*)^, respectively. Given these factor matrices, let the current estimate of the training tensor be

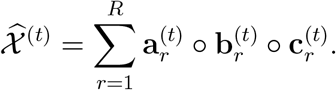

Define the element-wise squared difference between two order-three tensors 𝒳 ∈ ℝ^*I*×*J*×*K*^ and 𝒴 ∈ ℝ^*I*×*J*×*K*^ as

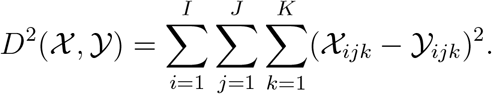

Alternating least squares on this tensor of order three is given by the following three steps:

1. Step 1: fix **A**^(*t*)^ and **B**^(*t*)^ and solve for **C**^(*t*+1^)

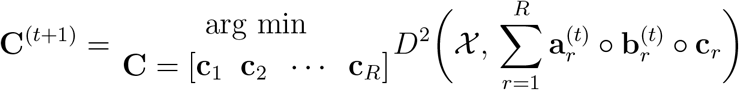
2. Step 2: fix **A**^(*t*)^ and **C**^(*t*)^ and solve for **B**^(*t*+1)^

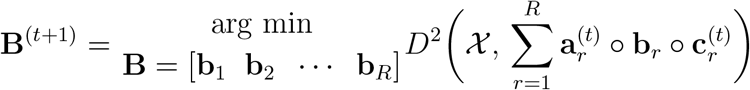
3. Step 3: fix **B**^(*t*)^ and **C**^(*t*)^ and solve for **A**^(*t*+1)^

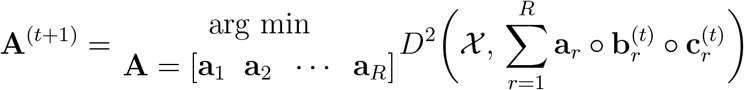

Steps 1 to 3 are repeated until convergence, and final estimated factor matrices are identical to those from the final iteration—*i*.*e*., **A** = **A**^(*t*+1)^, **B** = **B**^(*t*+1)^, and **C** = **C**^(*t*+1)^. At each step we incorporate the values for *λ*_*r*_, *r* = 1, 2, …, *R* into the estimated tensor.

#### Projecting test observations onto identified factor matrices

Given a new test tensor 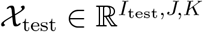 of *I*_test_ test observations, we can project the test observations onto the learned factor **A** so that it falls within the subspace learned by decomposing the training tensor 𝒳. However, before doing so, we must ensure that the test dataset lies in the same input space as the training set. Thus, we preprocess the test dataset by applying Equations 1 and 2 for centering and scaling. It is important to note that Equation 1 refers to centering with respect to the training set (*i*.*e*., subtracting 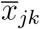), and so the test set must be centered with the mean training pixel value 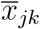 and not a similar quantity for the test set. Thus, the centering values for the training set must be retained so that the test set is centered with identical values. Assuming 𝒳_test_ has now been properly preprocessed, we can project the test data onto the learned features representing each input image by [Kolda and Bader, 2009]

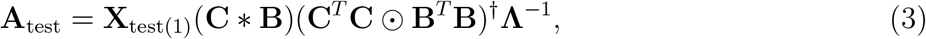

where **X**_test(1)_ is the mode-1 unfolding (matricization) of the tensor 𝒳_test_, the superscript *T* denotes transpose, the symbol ∗ denotes the Khatri-Rao product, the ⊙ symbol denotes the Hadamard (element-wise) product, the superscript † denotes the Moore-Penrose pseudoinverse, and **Λ**^*−*1^ represents the inverse of the diagonal matrix **Λ** ∈ ℝ^*R*×*R*^ of scaling terms *λ*_1_, *λ*_2_, …, *λ*_*R*_.

### Alignment processing

We used a novel approach for processing the haplotype alignments in a way that helps the classifiers detect the footprint of a selective sweep. For each simulated 1.1 Mb region, we locally sorted haplotypes in windows of 100 single nucleotide polymorphisms (SNPs), moving the window along the region with a stride of 10 SNPs, where values at SNPs were averaged for all windows that overlapped them. This method of alignment processing can help classifiers identify signals of lost haplotypic diversity if sweeps are weak or old, while also retaining power for strong and recent sweeps. To reduce the complexity of the tensor decomposition and noise in the sorted haplotype alignments, we downsampled the alignment images to 64×64-dimensional matrices using the `scikit-image` library [Pedregosa et al., 2011], where Gaussian smoothing was employed to preserve the spatial relationships of pixels within the images and to avoid aliasing artifacts. We highlight the advantage of our alignment processing approach by pitting the results obtained after employing our unique alignment processing strategy against those of an alignment processing approach that is similar to that used by `ImaGene` [Torada et al., 2019] (compare Figure S12 to Figure S1 and Figure S13 to Figure S2).

### *T-REx* model training and hyperparameter tuning

We have implemented three classical linear and nonlinear machine learning models with different R packages into our *T-REx* framework. For performing tensor decomposition, we used the R package `rTensor` [Li et al., 2018]. Additionally, we employed the R packages `glmnet` [Friedman et al.], `liquidsvm` [Steinwart and Thomann, 2017], and `ranger` [Wright and Ziegler, 2017] for implementing *T-REx*(EN), *T-REx*(SVM), and *T-REx*(RF), respectively. During the training of each classifier, we have 10^4^ observations in each class, with each observation consisting of sorted haplotype alignments (details provided in the *Alignment processing* subsection of the *Methods*). We then applied a rank *R* tensor decomposition (see *CP tensor decomposition* subsection of the *Methods* for details) to obtain a set of *R* derived features for each observation in each class to be used as input for our *T-REx* classifiers.

Before the testing phase commences, we tuned hyperparameters, which control certain components of the model training process, of each model by selecting optimal hyperparameters through the cross validation procedure. Hyperparameter tuning is a way of selecting suitable hyperparameter values from a range of possible values. Specifically, we performed 10-fold cross validation such that on each of the 10 folds we selected 10% of the samples (1,000 observations per class) from the dataset to be reserved for model validation and the remaining 90% of the samples (9,000 observations per class) to be employed for model training. This procedure allowed us to evaluate how well the model would perform on unseen data (from the validation set) for a given set of hyperparameter values. For each of the classifiers, we chose the model structure that yielded the smallest cross validation error after performing hyperparameter tuning.

For hyperparameter tuning of *T-REx*(EN), we explored a grid of values *α* ∈ {0, 0.1, …, 1.0}, where *α* denotes the proportion of the model for which the parameters are penalized with an *L*_2_-norm penalty, whereas 1 − *α* is the proportion penalized with an *L*_1_-norm penalty [Hastie et al., 2009]. In addition to *α*, we tuned another hyperparameter *λ* ≥ 0, which modulates the complexity of the fitted model by controlling the influence of the *L*_1_- and *L*_2_-norm penalties during model training. By tuning both *λ* and *α*, we are controlling the complexity of the fitted model while simultaneously performing feature selection by inclusion of the *L*_1_-norm penalty [Hastie et al., 2009]. We find the optimal *λ* and *α* combination that gives the minimum 10-fold cross validation error. For implementing *T-REx*(SVM), we used the radial basis kernel for nonlinear modeling, which has a hyperparameter *γ* that is inversely proportional to the variance (width) of the radial basis kernel, which has a shape similar to a Gaussian function [Hastie et al., 2009]. To implement *T-REx*(RF), we chose a large number (5,000) of random trees to use in the random forest ensemble, as test error stabilizes with enough trees in the forest [Hastie et al., 2009], and used the default number of 10 random splits within `ranger` for growing each decision tree within the random forest.

Finally, regardless of machine learning method, another important hyperparameter is the rank *R* of the tensor tensor decomposition. For each value of *R* ∈ {50, 100, 150, 200, 250, 300}, we computed the 10-fold cross validation error for *T-REx*(EN) and *T-REx*(SVM) and the out-of-bag error for *T-REx*(RF) [Hastie et al., 2009]. We chose the (*R, λ, α*) triple that resulted in the smallest 10-fold cross validation error for *T-REx*(EN), the (*R, γ*) pair that results in the smallest 10-fold cross validation for *T-REx*(SVM), and the value of *R* that results in the smallest out-of-bag error for *T-REx*(RF). After selecting the set of optimal hyperparameters of each method, the three *T-REx* models were each trained on the full dataset of 10^4^ training observations per class conditional on their optimal hyperparameters, and these models were deployed on further testing data.

### Training and evaluating ImaGene

To fully evaluate the performance *T-REx*, we compared it with the CNN-based sweep classifier `ImaGene` (details are provided in the *Results*). While both *T-REx* and `ImaGene` use haplotype alignments in the form of images, there are differences in the procedure used to process the images and perform model training. For training, `ImaGene` employs a “simulation-on-the-fly” approach of using newly generated data at each training epoch (iteration of gradient descent). This simulation-on-the-fly approach prevents `ImaGene` from overfitting. For consistency and fairness in comparison between *T-REx* and `ImaGene`, we deviated from this default setting of `ImaGene` so that it is pitted against *T-REx* on identical simulation data. Specifically, we used the same 10^4^ training observations per class when training `ImaGene` as we employed for training *T-REx* for each simulation setting (details regarding the simulation protocol are provided in the *Results*). To prevent overfitting, we employed early stopping [Goodfellow et al., 2016a], by setting the number of epochs to train `ImaGene` as the point at which the validation loss starts to rise, which suggests overfitting, where the validation loss was computed across 1,000 observations per class that were held out for validation. Figure S14 displays the validation and training loss curves over 200 training epochs, showing that the validation curve begins to increase at approximately 25 epochs. We therefore retrained the `ImaGene` model on the full dataset of 10^4^ observations per class for 25 epochs.

### Application to empirical data

With the aim of detecting novel candidate genes that may be subject to positive natural selection and previously hypothesized candidates of positive natural selection, we used empirical data of the central European human population CEU from the 1000 Genomes Project dataset [The 1000 Genomes Project Consortium, 2015]. We first filtered variant calls to include biallelic SNPs. Second, we removed SNPs with minor allele count less than three, as Mughal et al. [2020] demonstrated the frequencies of singleton and doubleton SNPs in the CEU population from the 1000 Genomes Project dataset differed from those predicted by the inferred demographic model [Terhorst et al., 2019] that we used to train our classifiers. Moreover, because regions of the genome that are harder to map and align may lead to technical artifacts affecting observed genomic variation [Derrien et al., 2012], we removed sites that could have problematic mapping or alignability to circumvent such potential artifacts. Specifically, we used the CRG score to measure mappability and alignability of a genomic region and removed sites falling within 100 kb windows for which the mean CRG100 score within the window was less than 0.9 [Mughal et al., 2020]. We then applied our unique alignment processing approach to further process the data before supplying it to *T-REx*.

## Acknowledgments

This work was supported by National Institutes of Health grant R35GM128590 and by National Science Foundation grants DEB-1949268, BCS-2001063, and DBI-2130666. Computations for this research were performed using the services provided by Research Computing at the Florida Atlantic University.

