## Supplemental figures for "Tensor decomposition based feature extraction and classification to detect natural selection from genomic data"

### Supplementary material

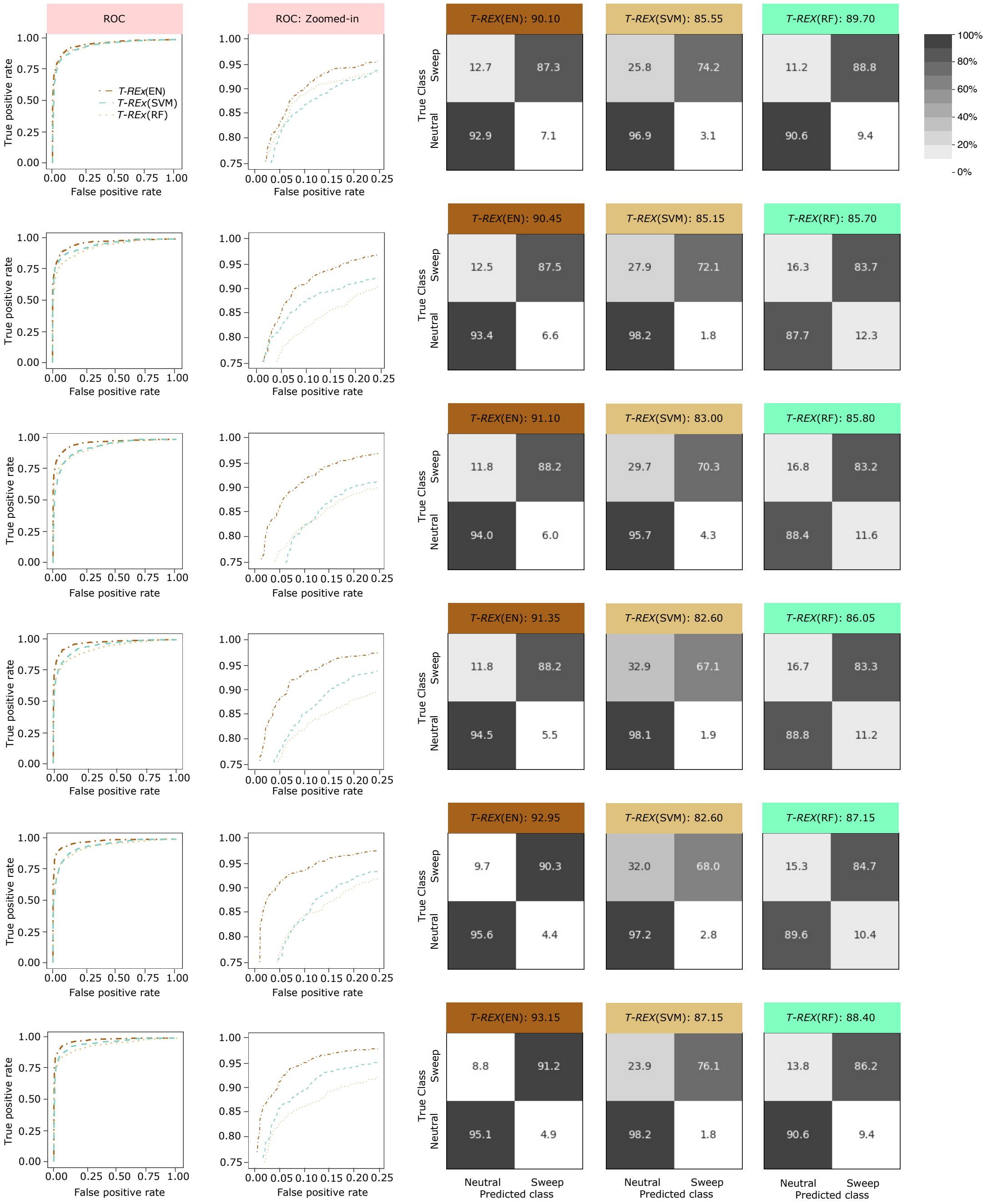


Figure S1: Powers and accuracies to detect sweeps for the linear *T-REx*(EN) and nonlinear *T-REx*(SVM) and *T-REx*(RF) classifiers applied to the `Constant_1` dataset for $R = 50$, 100, 150, 200, 250, and 300 (top row to bottom), respectively. For training and testing purposes, the number of observations used for each class was 10,000 and 1000, respectively. Powers to detect sweeps of all three methods are compared using receiver operating characteristic (ROC) curves (first column) and ROC curves zoomed in to the upper left-hand corner with false positive rate less than 0.25 and true positive rate greater than 0.75 (second column). Classification accuracy and rates of all three methods are depicted using confusion matrices in columns three through five for *T-REx*(EN), *T-REx*(SVM), and *T-REx*(RF), respectively.

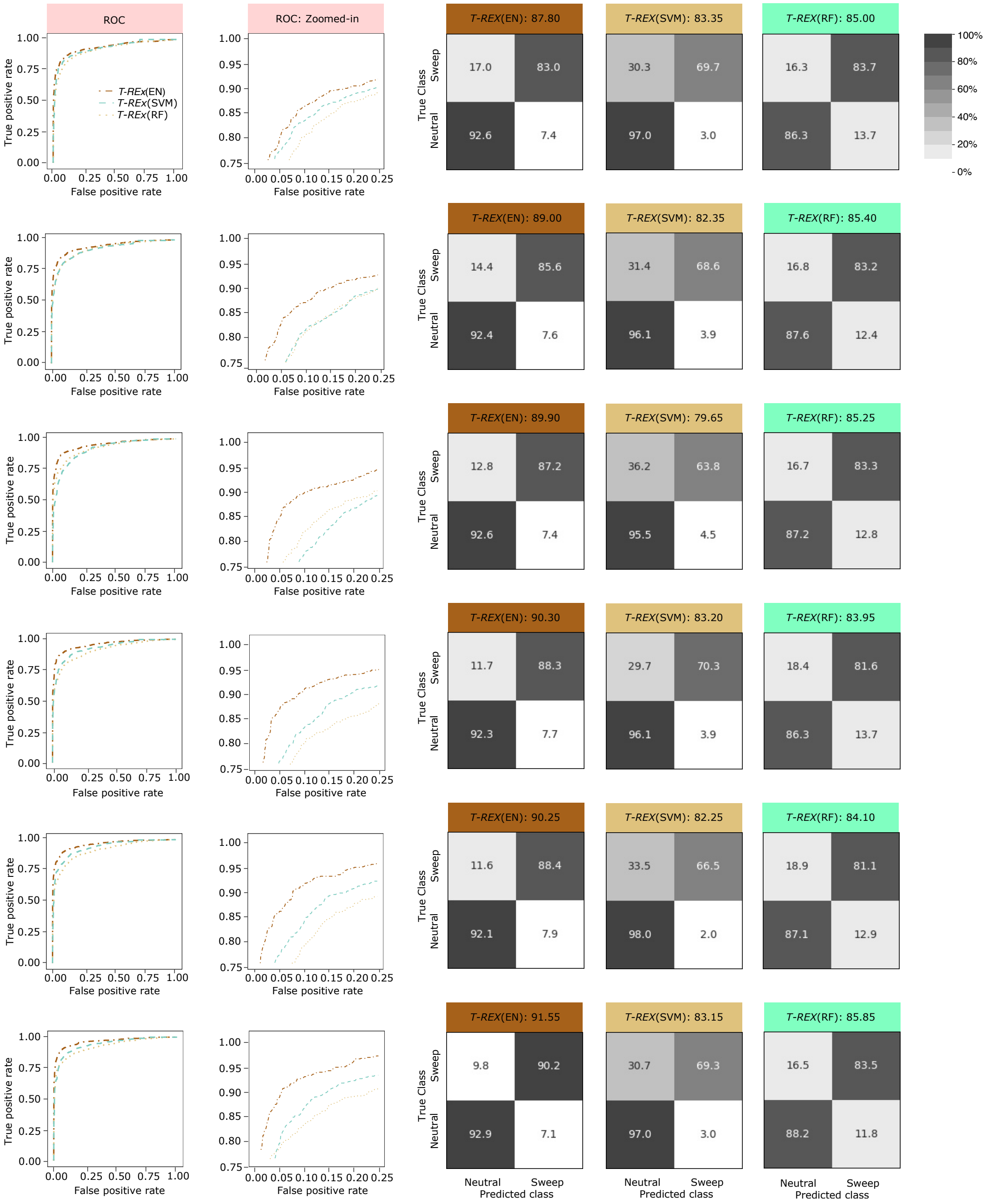


Figure S2: Powers and accuracies to detect sweeps for the linear *T-REx*(EN) and nonlinear *T-REx*(SVM) and *T-REx*(RF) classifiers applied to the `Constant_2` dataset for $R$ = 50, 100, 150, 200, 250, and 300 (top row to bottom), respectively. For training and testing purposes, the number of observations used for each class was 10,000 and 1000, respectively. Powers to detect sweeps of all three methods are compared using receiver operating characteristic (ROC) curves (first column) and ROC curves zoomed in to the upper left-hand corner with false positive rate less than 0.25 and true positive rate greater than 0.75 (second column). Classification accuracy and rates of all three methods are depicted using confusion matrices in columns three through five for *T-REx*(EN), *T-REx*(SVM), and *T-REx*(RF), respectively.

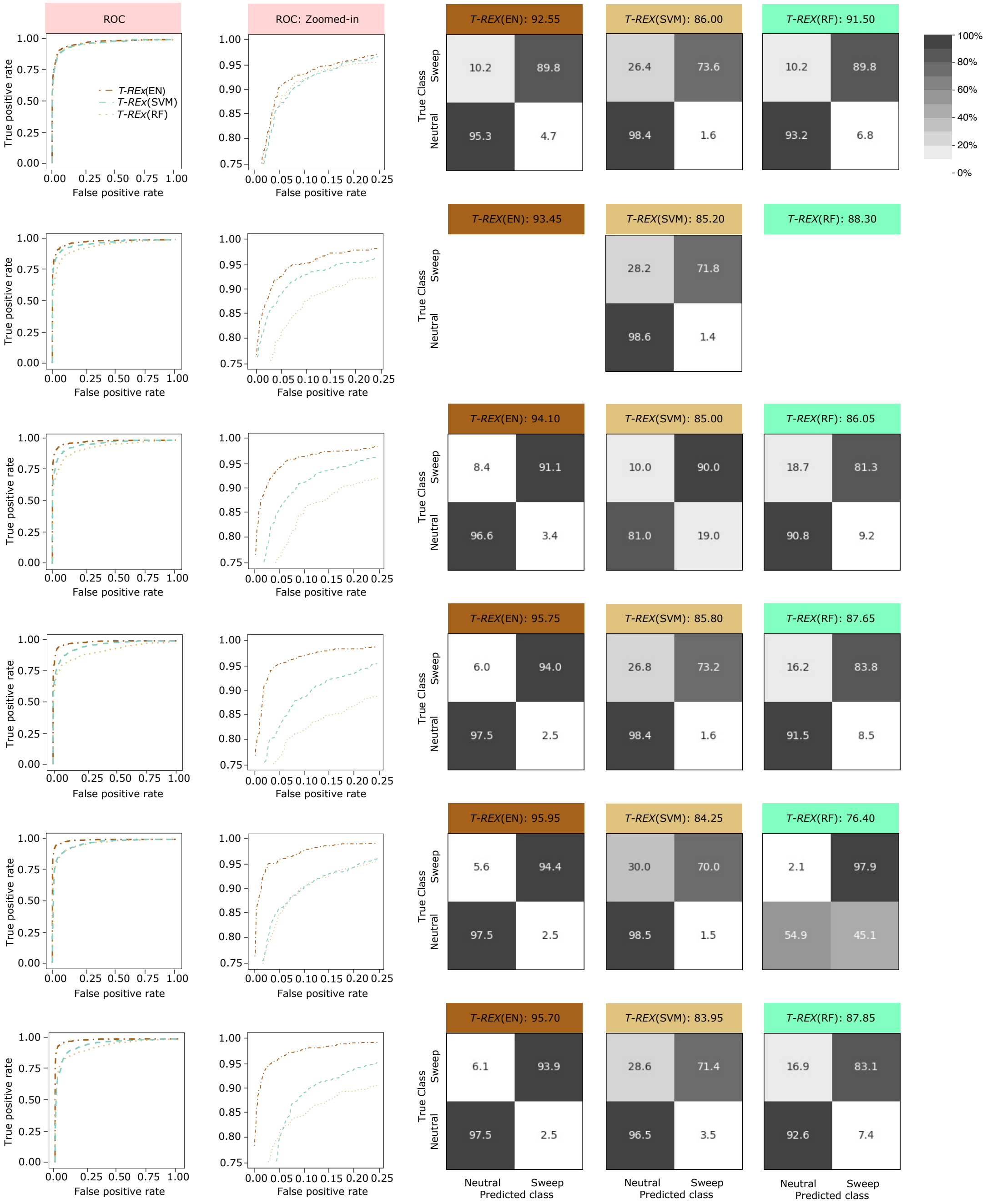


Figure S3: Powers and accuracies to detect sweeps for the linear *T-REx*(EN) and nonlinear *T-REx*(SVM) and *T-REx*(RF) classifiers applied to the CEU_1 dataset for $R$ = 50, 100, 150, 200, 250, and 300 (top row to bottom), respectively. For training and testing purposes, the number of observations used for each class was 10,000 and 1000, respectively. Powers to detect sweeps of all three methods are compared using receiver operating characteristic (ROC) curves (first column) and ROC curves zoomed in to the upper left-hand corner with false positive rate less than 0.25 and true positive rate greater than 0.75 (second column). Classification accuracy and rates of all three methods are depicted using confusion matrices in columns three through five for *T-REx*(EN), *T-REx*(SVM), and *T-REx*(RF), respectively.

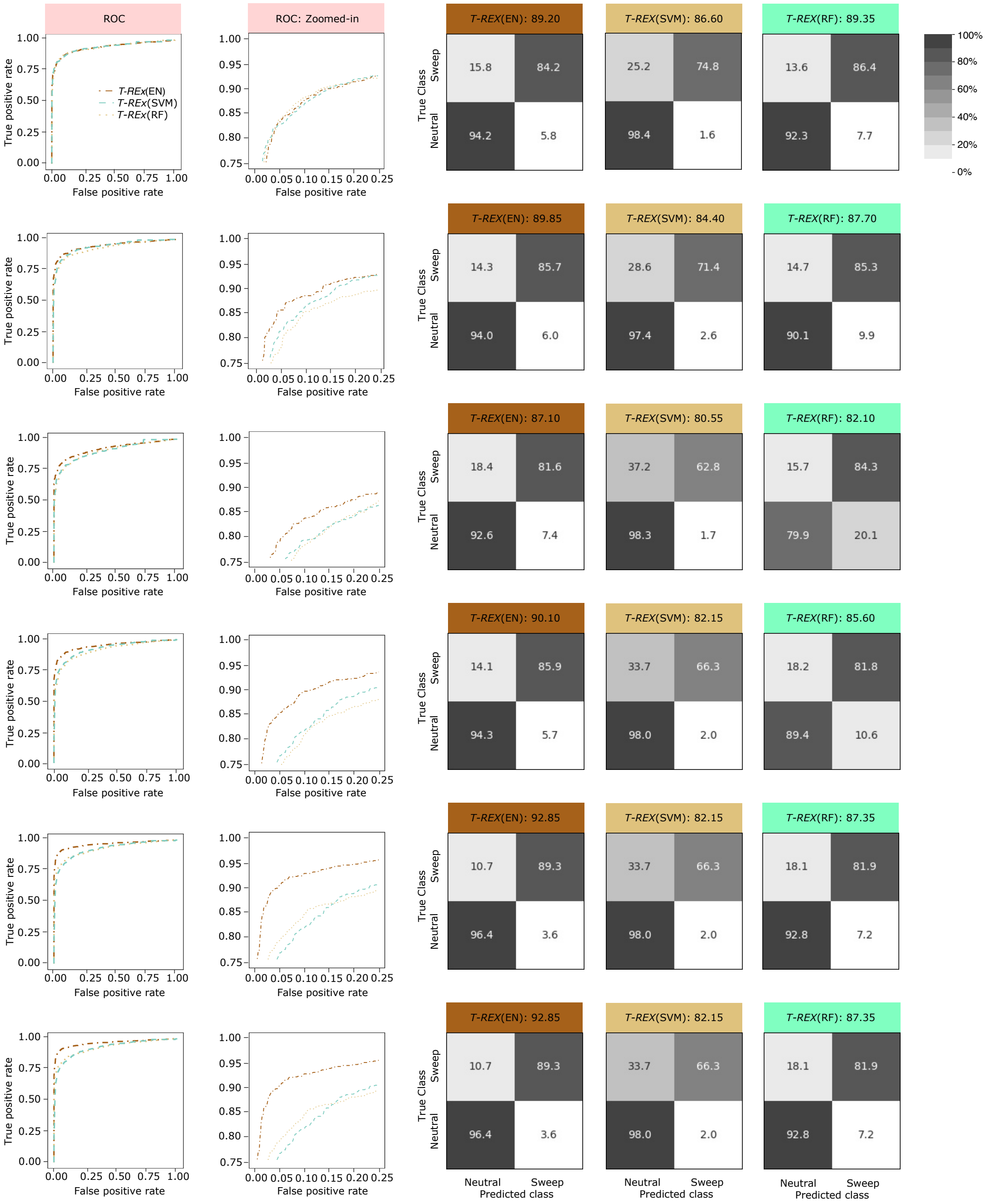


Figure S4: Powers and accuracies to detect sweeps for the linear *T-REx*(EN) and nonlinear *T-REx*(SVM) and *T-REx*(RF) classifiers applied to the `CEU_2` dataset for $R$ = 50, 100, 150, 200, 250, and 300 (top row to bottom), respectively. For training and testing purposes, the number of observations used for each class was 10,000 and 1000, respectively. Powers to detect sweeps of all three methods are compared using receiver operating characteristic (ROC) curves (first column) and ROC curves zoomed in to the upper left-hand corner with false positive rate less than 0.25 and true positive rate greater than 0.75 (second column). Classification accuracy and rates of all three methods are depicted using confusion matrices in columns three through five for *T-REx*(EN), *T-REx*(SVM), and *T-REx*(RF), respectively.

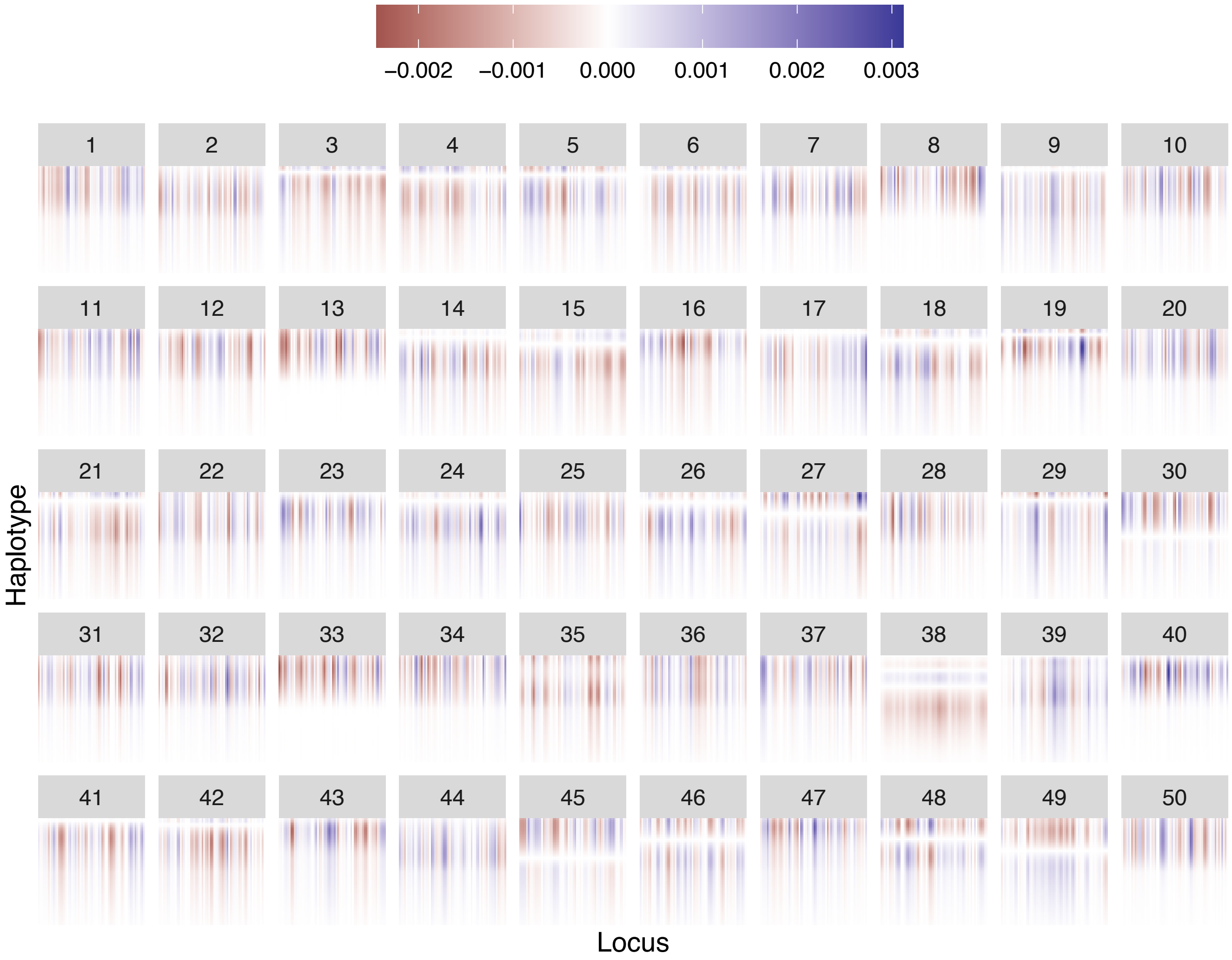


Figure S5: Feature maps visualizing the latent structure of the training data after performing CP decomposition. The resultant $R$ components have been sorted such that $\lambda_1 \geq \lambda_2 \geq \cdots \geq \lambda_R$, where $\lambda_r$ is the $r$th element in the diagonal of the core tensor used in CP decomposition. Sorted components 1 through 50, out of $R = 250$ components extracted from the `CEU_2` dataset are depicted.

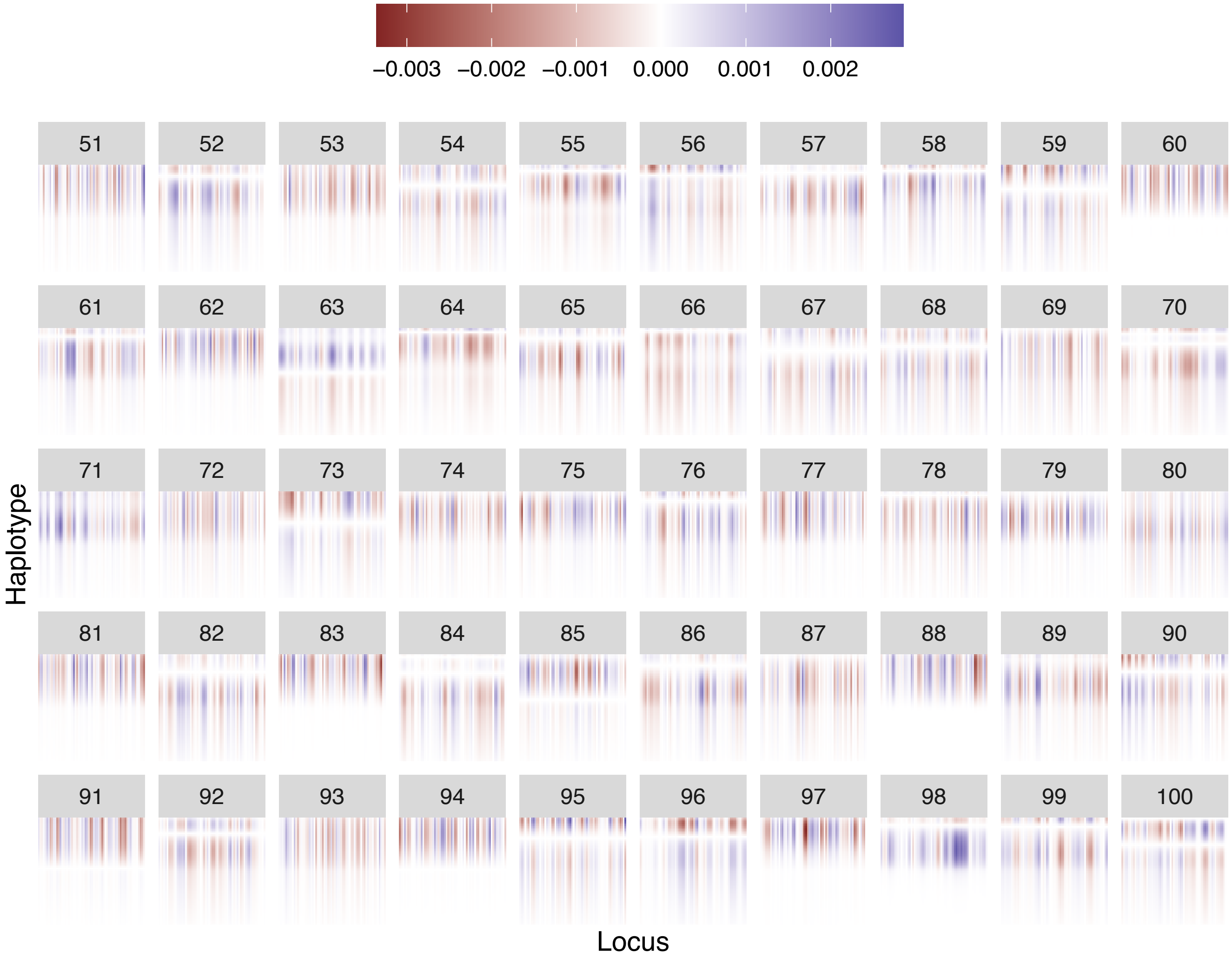


Figure S6: Feature maps visualizing the latent structure of the training data after performing CP decomposition. The resultant $R$ components have been sorted such that $\lambda_1 \geq \lambda_2 \geq \cdots \geq \lambda_R$, where $\lambda_r$ is the $r$th element in the diagonal of the core tensor used in CP decomposition. Sorted components 51 through 100, out of $R = 250$ components extracted from the `CEU_2` dataset are depicted.

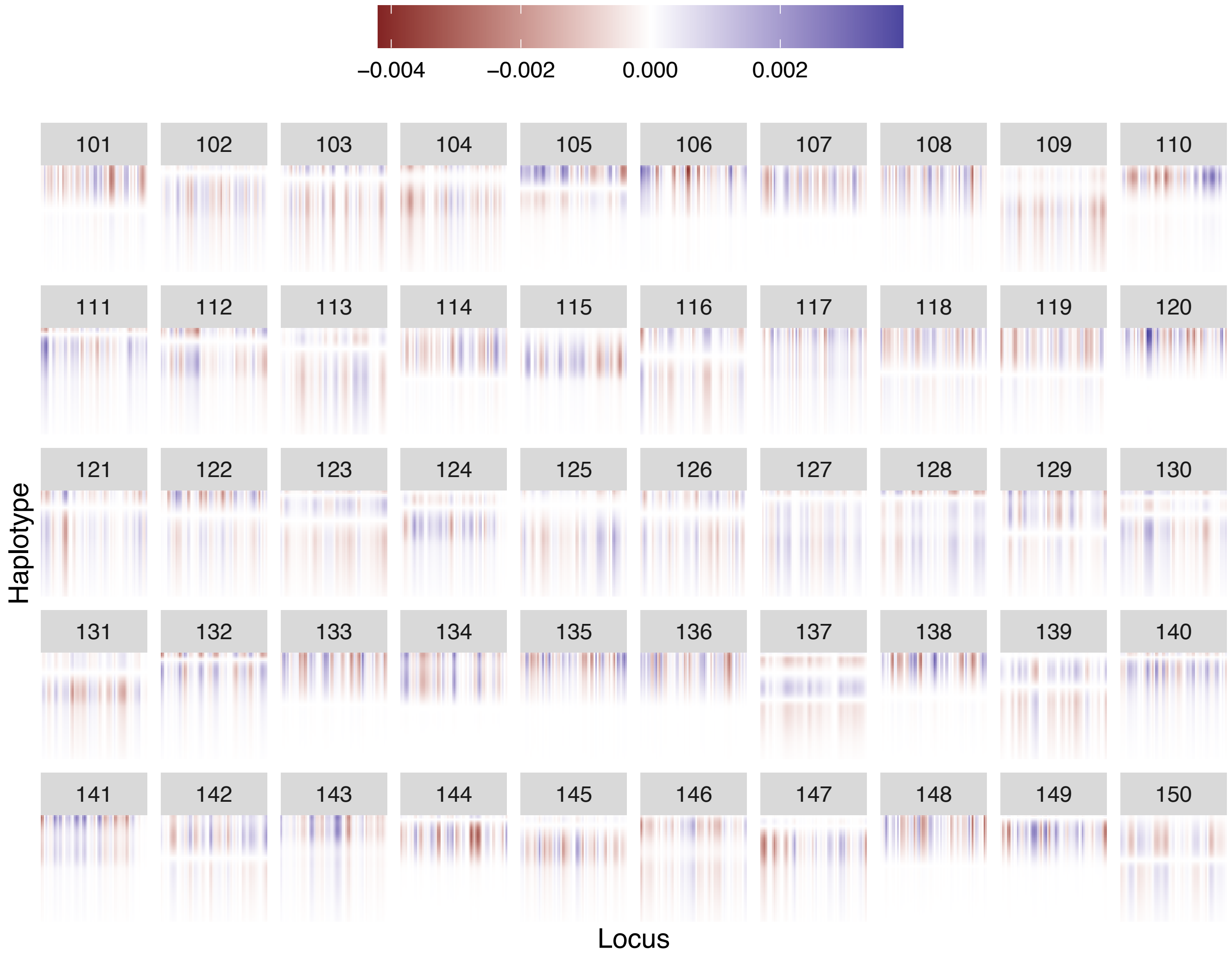


Figure S7: Feature maps visualizing the latent structure of the training data after performing CP decomposition. The resultant $R$ components have been sorted such that $\lambda_1 \geq \lambda_2 \geq \cdots \geq \lambda_R$, where $\lambda_r$ is the $r$th element in the diagonal of the core tensor used in CP decomposition. Sorted components 101 through 150, out of $R = 250$ components extracted from the CEU_2 dataset are depicted.

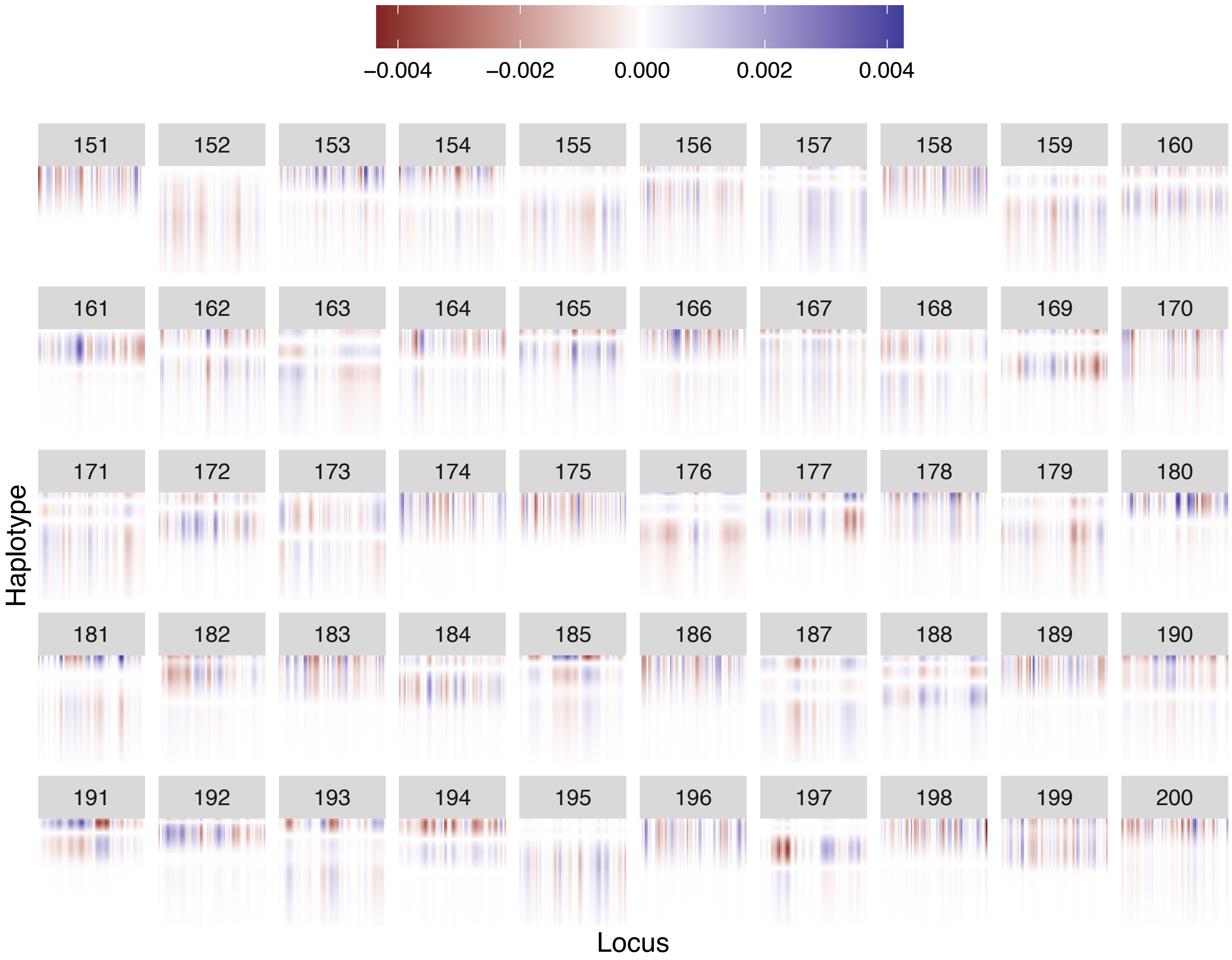


Figure S8: Feature maps visualizing the latent structure of the training data after performing CP decomposition. The resultant $R$ components have been sorted such that $\lambda_1 \geq \lambda_2 \geq \cdots \geq \lambda_R$, where $\lambda_r$ is the $r$th element in the diagonal of the core tensor used in CP decomposition. Sorted components 151 through 200, out of $R = 250$ components extracted from the CEU_2 dataset are depicted.

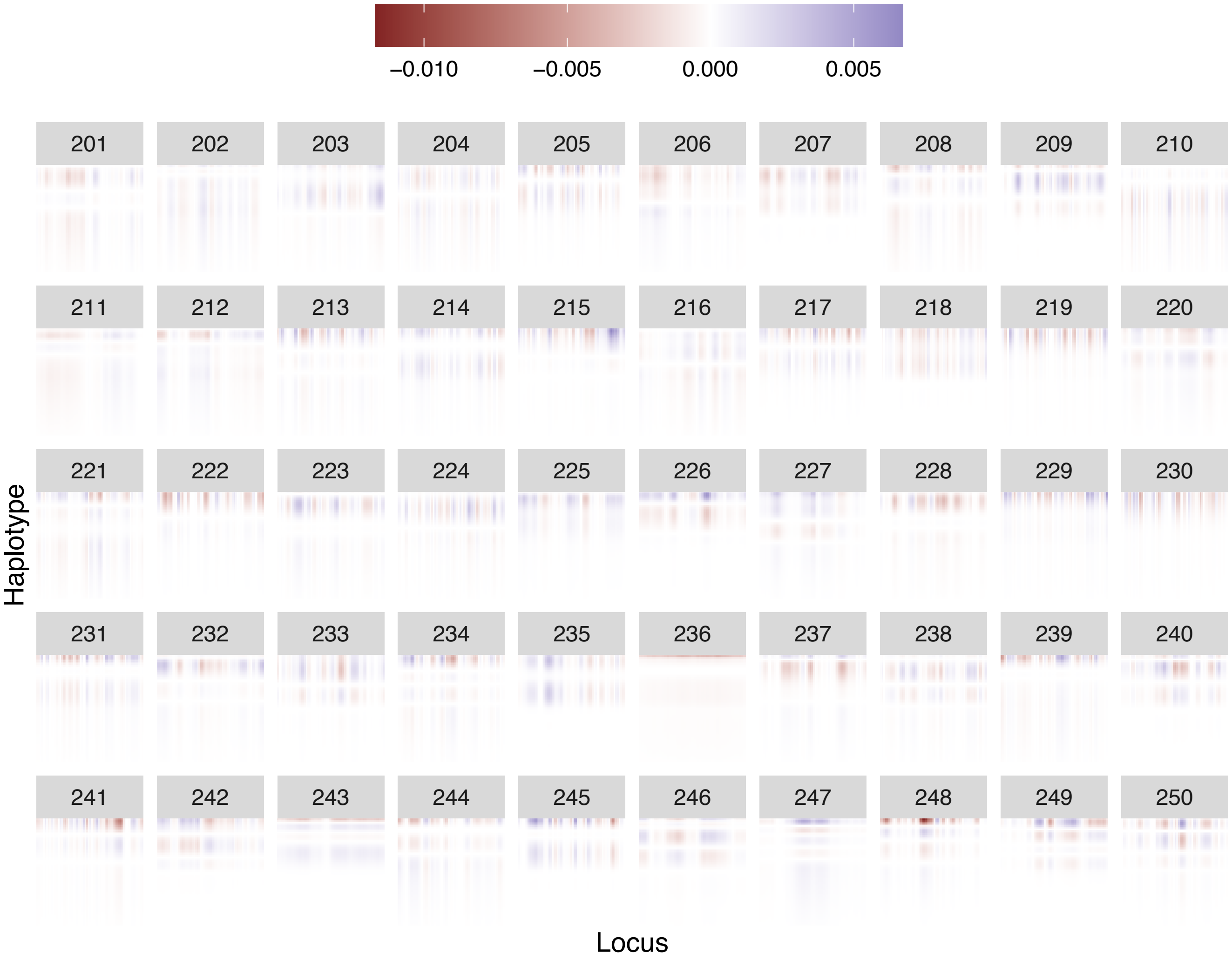


Figure S9: Feature maps visualizing the latent structure of the training data after performing CP decomposition. The resultant $R$ components have been sorted such that $\lambda_1 \geq \lambda_2 \geq \cdots \geq \lambda_R$, where $\lambda_r$ is the $r$th element in the diagonal of the core tensor used in CP decomposition. Sorted components 201 through 250, out of $R = 250$ components extracted from the `CEU_2` dataset are depicted.

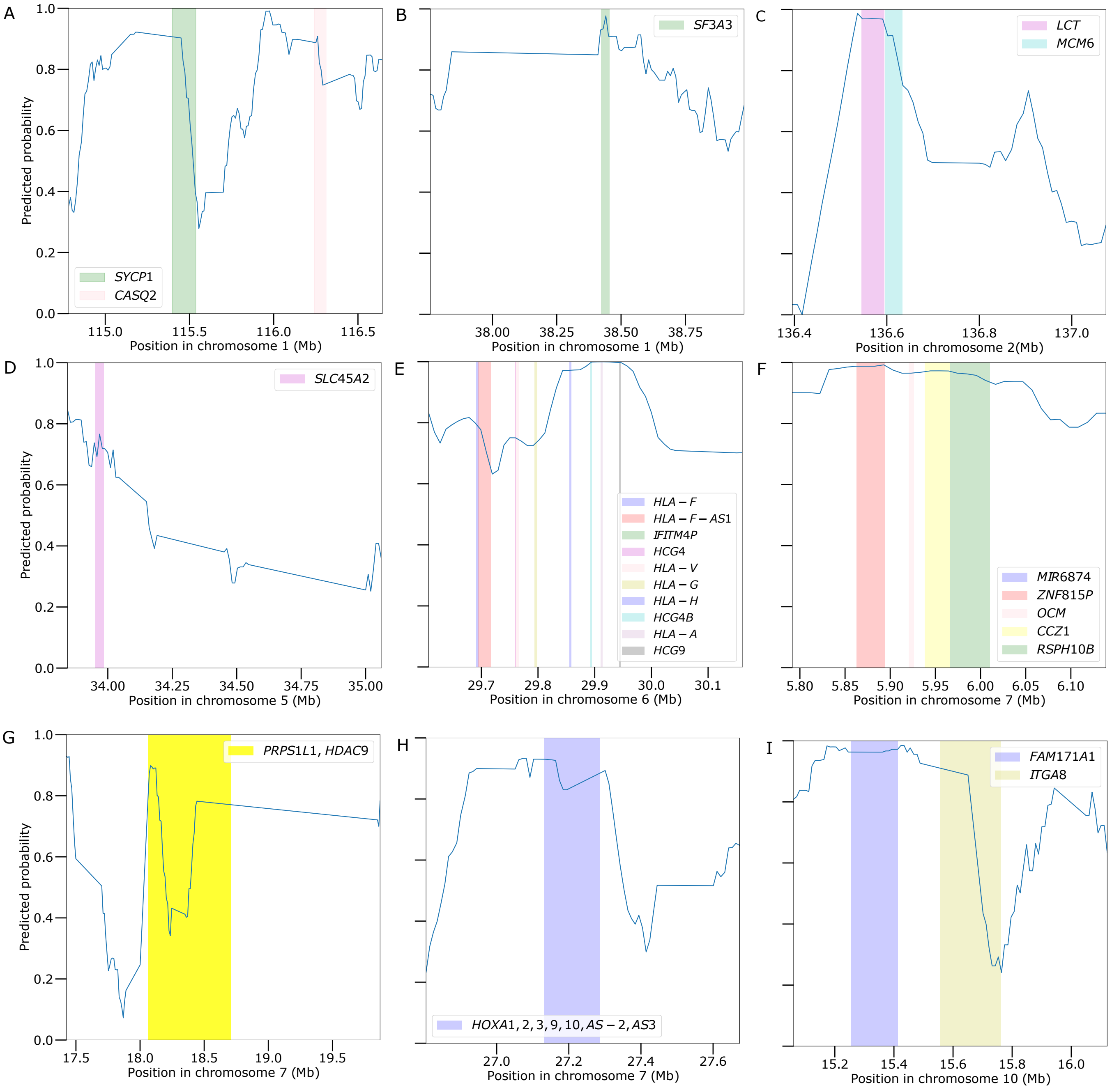


Figure S10: Sweep probabilities predicted by *T-REx*(EN) surrounding regions of interest on chromosomes 1, 2, 5, 6, 7, and 10 of the CEU population as a function of chromosomal position. The blue smoothed curve is capturing the eleven point moving average, computed with five windows before and five windows after a given central window. The genomic intervals containing each gene are shaded using colors in accordance with the order of their appearance in the labels.

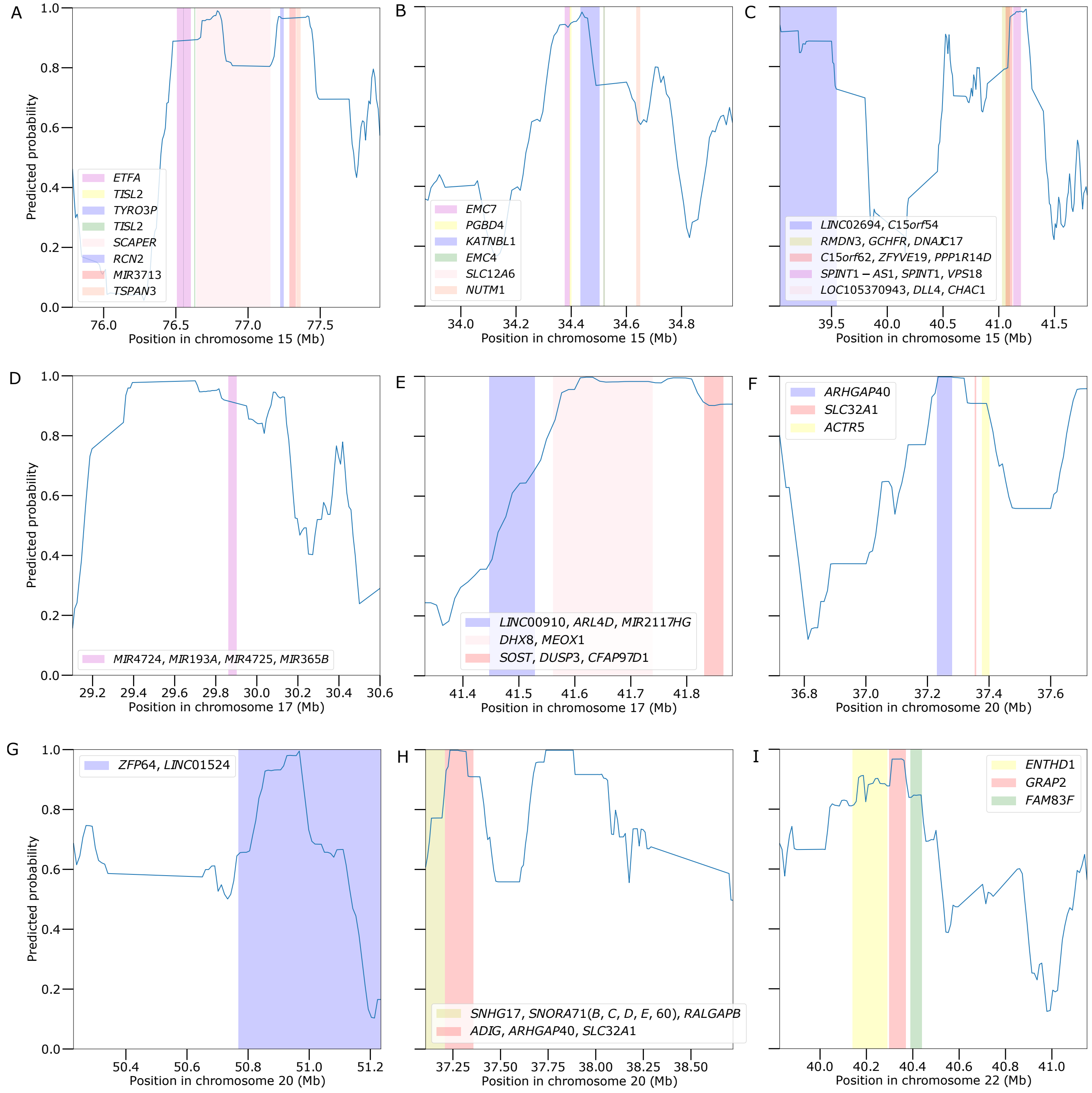


Figure S11: Sweep probabilities predicted by $T$-$REx$(EN) surrounding regions of interest on chromosomes 15, 17, 20, and 22 of the CEU population as a function of chromosomal position. The blue smoothed curve is capturing the eleven point moving average, computed with five windows before and five windows after a given central window. The genomic intervals containing each gene are shaded using colors in accordance with the order of their appearance in the labels.

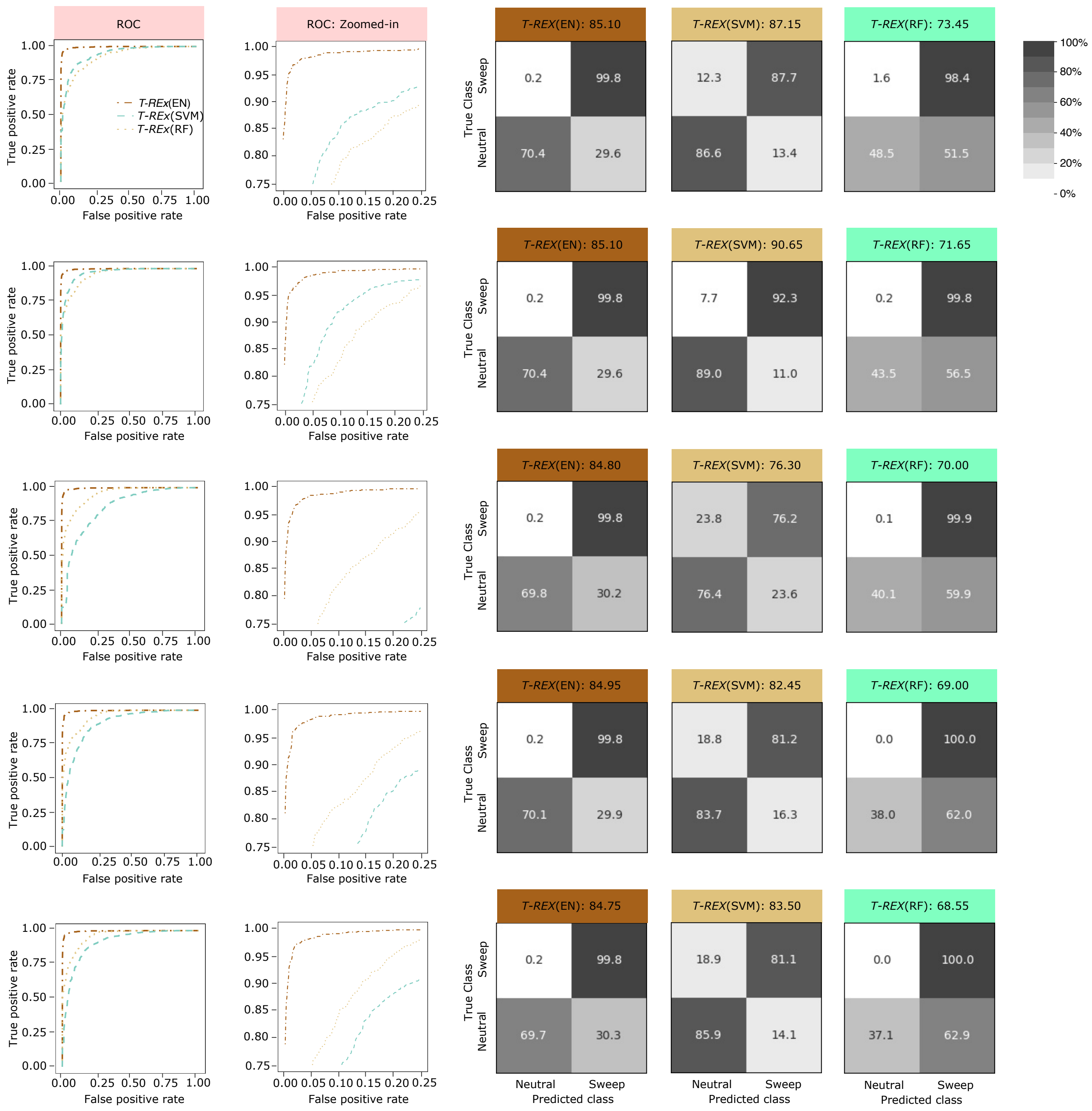


Figure S12: Powers and accuracies to detect sweeps for the linear *T-REx*(EN) and nonlinear *T-REx*(SVM) and *T-REx*(RF) classifiers applied to the `Constant_1` dataset using an alignment processing approach that was similar to the one used by Torada et al. [2019] for $R$ = 50, 100, 150, 200, and 250 (top row to bottom), respectively. For training and testing purposes, the number of observations used for each class was 10,000 and 1000, respectively. Powers to detect sweeps of all three methods are compared using receiver operating characteristic (ROC) curves (first column) and ROC curves zoomed in to the upper left-hand corner with false positive rate less than 0.25 and true positive rate greater than 0.75 (second column). Classification accuracy and rates of all three methods are depicted using confusion matrices in columns three through five for *T-REx*(EN), *T-REx*(SVM), and *T-REx*(RF), respectively.

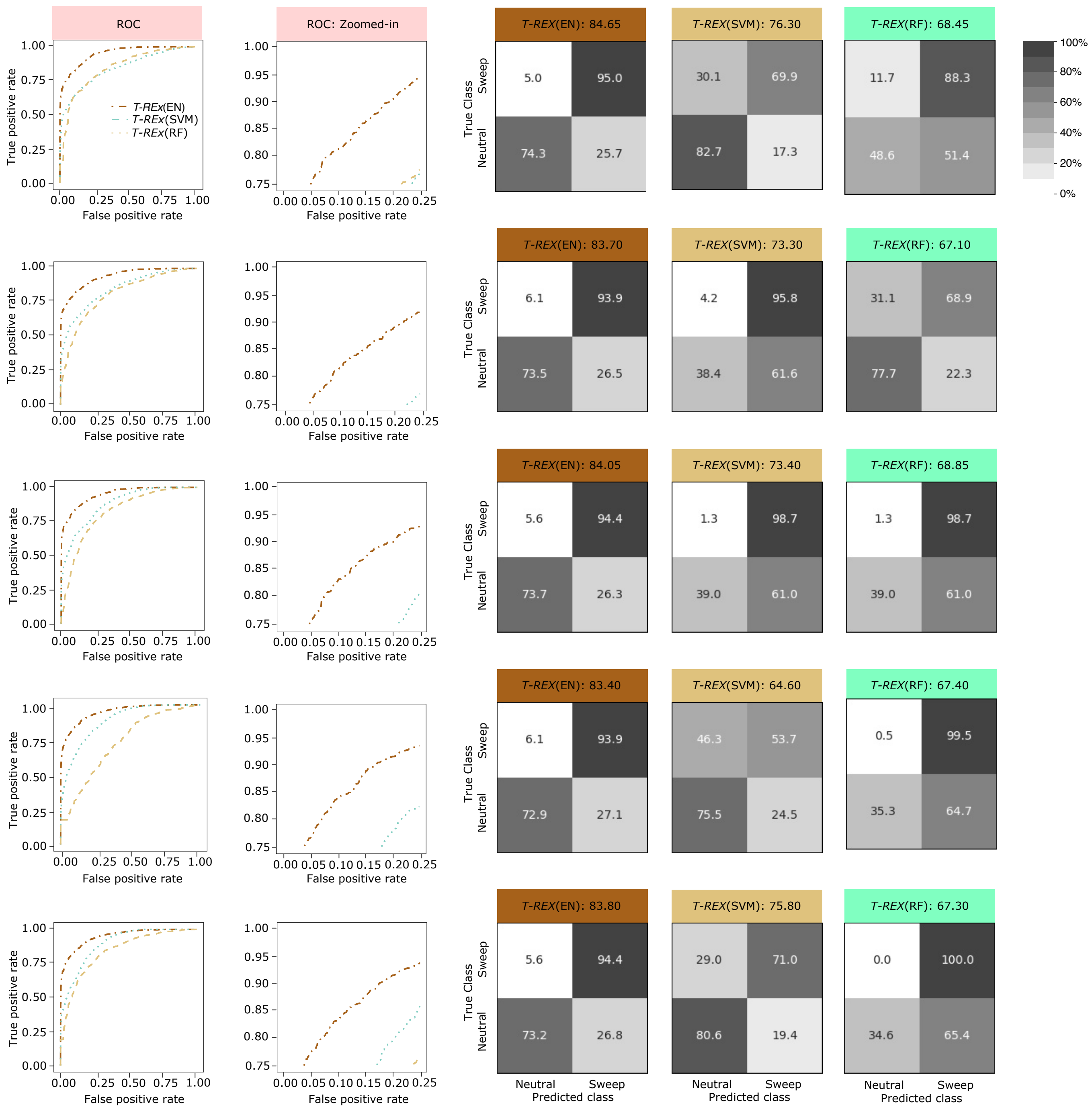


Figure S13: Powers and accuracies to detect sweeps for the linear *T-REx*(EN) and nonlinear *T-REx*(SVM) and *T-REx*(RF) classifiers applied to the `Constant_2` dataset using an alignment processing approach that was similar to the one used by Torada et al. [2019] for $R$ = 50, 100, 150, 200, and 250 (top row to bottom), respectively. For training and testing purposes, the number of observations used for each class was 10,000 and 1000, respectively. Powers to detect sweeps of all three methods are compared using receiver operating characteristic (ROC) curves (first column) and ROC curves zoomed in to the upper left-hand corner with false positive rate less than 0.25 and true positive rate greater than 0.75 (second column). Classification accuracy and rates of all three methods are depicted using confusion matrices in columns three through five for *T-REx*(EN), *T-REx*(SVM), and *T-REx*(RF), respectively.

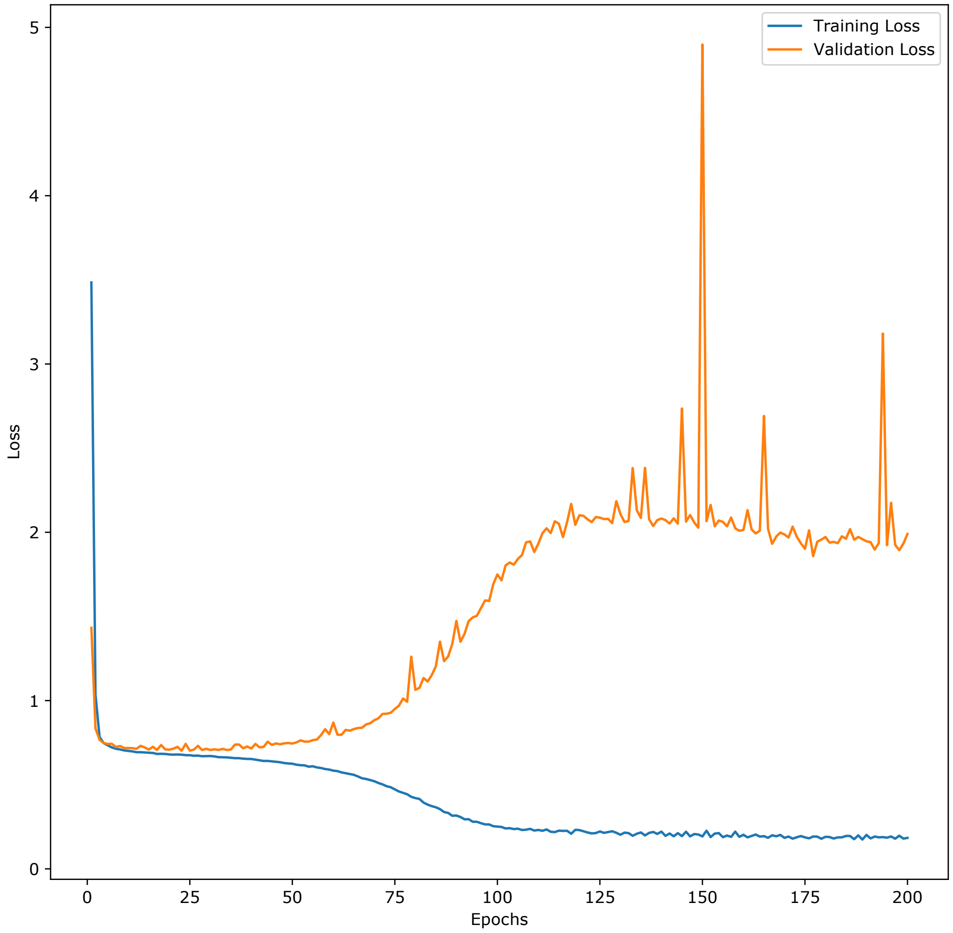


Figure S14: `ImaGene` training and validation loss curves as a function number of training epochs. Due to the continuous increase of the validation loss after 25 epochs, we selected to employ 25 epochs to train the final `ImaGene` model to avoid overfitting.
